# Structure of the human FERRY Rab5 effector complex

**DOI:** 10.1101/2021.06.21.449265

**Authors:** Dennis Quentin, Jan S. Schuhmacher, Björn U. Klink, Jeni Lauer, Tanvir R. Shaikh, Pim J. Huis in ’t Veld, Luisa M. Welp, Henning Urlaub, Marino Zerial, Stefan Raunser

## Abstract

Long-range mRNA transport is crucial for the spatio-temporal regulation of gene expression, and its malfunction leads to neurological disorders. The pentameric FERRY Rab5 effector complex is the molecular link between mRNA and early endosomes in mRNA intracellular distribution. Here, we determine the cryo-EM structure of the human FERRY complex, composed of Fy-1 to Fy-5. The structure reveals a clamp-like architecture, where two arm-like appendages of Fy-2 and a Fy-5 dimer, protrude from the central Fy-4 dimer. The coiled-coil domains of Fy-2 are flexible and project into opposite directions from the core. While the Fy-2 C-terminal coiled-coil acts as binding region for Fy-1/3 and Rab5, both coiled-coils and Fy-5 concur to bind mRNA. Mutations causing truncations of Fy-2 in patients with neurological disorders impair Rab5 binding or FERRY complex assembly. Thus, Fy-2 serves as a binding hub connecting all five complex subunits and mediating the binding to mRNA and early endosomes via Rab5. The FERRY structure provides novel mechanistic insights into long-distance mRNA transport.

## Introduction

Rab small GTPases are master regulators of membrane organelles (Pfeffer, 2017; Wandinger-Ness and Zerial, 2014; Zerial and McBride, 2001). They spatially and temporally regulate essential vesicular transport processes including organelle biogenesis, protein secretion, receptor internalization, recycling of membrane-associated molecules and cell-type specific trafficking. Thus, they contribute to the structural and functional integrity of organelles, which is critical for cellular homeostasis in eukaryotes.

In the GTP-bound conformation, membrane-associated Rab proteins can recruit a plethora of diverse downstream effector proteins to accomplish membrane remodeling activities (Grosshans et al.; Pfeffer, 2017; Wandinger-Ness and Zerial, 2014; Zerial and McBride, 2001).

Rab5, one of the most extensively studied Rab GTPases, is mainly localized at the early endosome (EE), and regulates endocytosis and EE dynamics (Woodman, 2000). A vast network of interaction partners of Rab5 has been identified, providing Rab5 with one of the most complex interactomes among the Rab family (Christoforidis et al., 1999). This includes GEFs like Rabex-5 and RIN1 (Horiuchi et al., 1997; Tall et al., 2001), but also Rab5-specific GAPs such as RN-Tre (Lanzetti et al., 2000) and Rab-GAP5 (Haas et al., 2005). Prominent effectors like Rabaptin-5, Rabankyrin-5, Rabenosyn-5 and APPL1/2 act downstream and can bind Rab5 via distinct domains such as zinc fingers (Miaczynska et al., 2004; Nielsen et al., 2000; Schnatwinkel et al., 2004; Stenmark et al., 1995, 1996). Similar to various members of the Ras superfamily, co-structures of Rab effectors bound to their cognate GTPase, including Rab5-Rabaptin-5, Rab4-Rabenosyn-5 and Rab11-FIP2, have provided important insights in the functional interactions between Rab proteins and their effectors and regulators (Eathiraj et al., 2005; Jagoe et al., 2006; Zhu et al., 2004). These studies demonstrate that binding is typically mediated by the switch and inter-switch regions of Rab proteins and either symmetric coiled-coils or α-helical bundles of effectors. In addition to individual proteins, Rab effectors comprise large multiprotein complexes that mediate crucial functions in the exocytic and endocytic pathways. These include the Exocyst and TRAPP, both involved in exocytosis (Sacher et al., 2001; Wu et al., 2010), and the two homologous hetero-hexameric tethering complexes, CORVET and HOPS, which interact with Rab5 and Rab7, respectively (Balderhaar and Ungermann, 2013; Kuhlee et al., 2015). All such large multiprotein complexes are non-symmetric, highly flexible and dynamic, and therefore challenging to analyze structurally. Hence, known structures are often limited to the core of the complexes, and it is difficult to reach high resolution (Bröcker et al., 2012; Chou et al., 2016; Kim et al., 2006).

Rab5 is also implicated in long-range endosomal motility (Hoepfner et al., 2005). By harnessing the intracellular microtubule (MT) network, EEs can be actively transported via MT motor complexes to distal locations within the cell. These long-distance transport processes are particularly important for cells with sophisticated internal architectures, such as neurons (Goto-Silva et al., 2019), explaining why deficiencies often manifest in compromised cognitive abilities.

Besides the spatial requirements of long-distance vesicular transport in exocytosis and endocytosis, RNA transport and local translation serves as a prime example of how spatio-temporal control can influence the expression of genes underlying essential biological processes, such as embryonic development or neuronal plasticity (Medioni et al., 2012; Mofatteh, 2020). Local sites are able to individually regulate gene expression, which is thus not limited to transcriptional control in the nucleus. The sophisticated mRNA localization pattern observed in polarized cells like neurons requires active transport of transcripts. Studies in a number of model systems, including yeast and *Drosophila melanogaster*, were performed in recent years to identify the relevant RNA transport machinery (Vazquez-Pianzola and Suter, 2012). Two distinct active transport pathways, both exploiting the cytoskeletal network in combination with motor proteins, have emerged: RNA is either transported with the help of accessory proteins, forming ribonucleoprotein particles (RNPs), or associated with endosomal compartments, both actively transported by co-opting cytoskeletal components. The role of late endosomes and lysosomes in mRNA localization has been subject of recent research, where Annexin 11A has been proposed to mediate the association between RNA and lysosomes (Liao et al., 2019). While these initial insights are valuable, they are limited to a subclass of the endocytic system. In filamentous fungi, mRNA localization is mediated by the microtubule-based long-distance transport of vesicles, including early endosomes (Zarnack and Feldbrügge, 2010). In neurons, long range transport of various types of cargo, including mRNA, requires active transport, which is mediated by endocytic organelles, particularly late endosomes (Cioni et al., 2019; Vos and Hafezparast, 2017). To date, little is known whether and how RNA is transported via early endosomes to its target destination.

We have identified a novel human 5-subunit Rab5 effector complex termed FERRY (Five-subunit Early endosome RNA and Ribosome intermediarY) complex (Schuhmacher et al., 2022), which interacts with mRNA and thus represents a prime candidate for early endosome-mediated mRNA transport. The FERRY complex is composed of five subunits Tbck (Fy-1), Ppp1r21 (Fy-2), C12orf4 (Fy-3), Cryzl1 (Fy-4), and Gatd1 (Fy-5), which have a molecular weight of 101, 88, 64, 39 and 23 kDa, respectively (Fig. 1A). Here, we determined the cryo-EM structure of the FERRY complex at a resolution of 4 Å. Together with rotary shadowing EM, hydrogen-deuterium exchange mass spectrometry (HDX-MS), crosslinking mass spectrometry (MS), electrophoretic mobility shift assay (EMSA) and mutational studies, the structure demonstrates that FERRY is an elongated complex with a clamp-like architecture at its center and protruding flexible coiled-coil structures at its periphery that mediate the interaction with the EE via Rab5 and mRNA.

**Figure 1.**
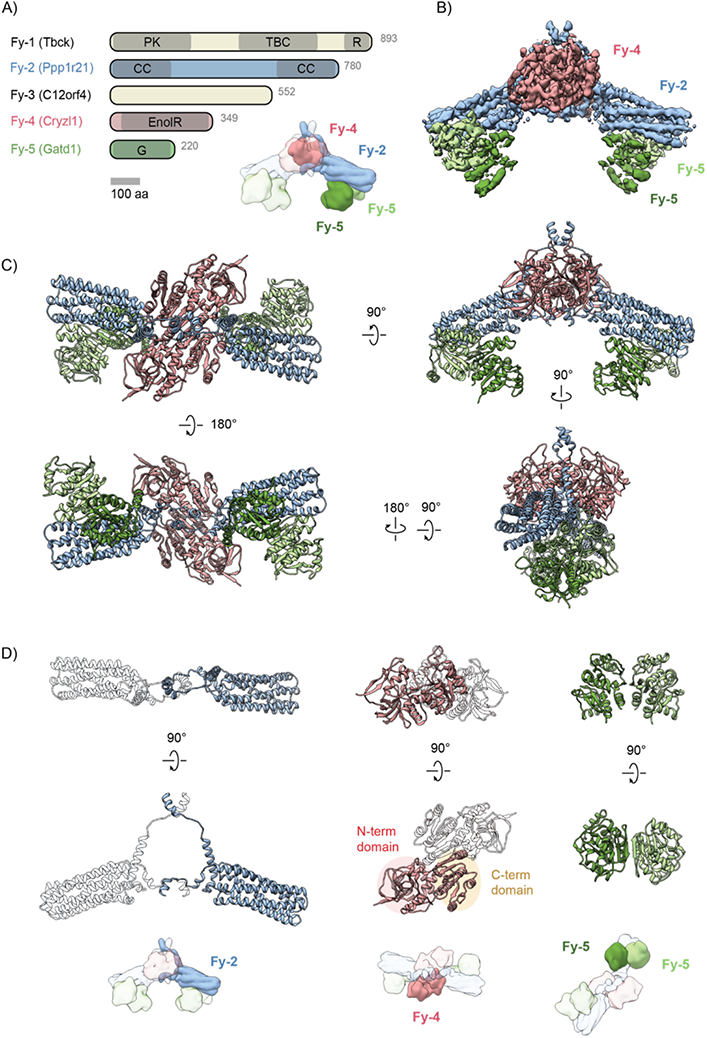
Architecture of the FERRY complex. (**A**) Domain architecture of the individual subunits of the FERRY complex (top, left). A schematic representation of the FERRY core is shown on the bottom right with only one half of the symmetric complex highlighted. Note that the asymmetric unit of the complex consists of one molecule of Fy-2 and Fy-4 as well as a dimer of Fy-5. Domain abbreviations: PK – Pseudokinase; TBC – Tre-2/Bub2/Cdc16; R – Rhodanese; CC – Coiled coil; EnolR – Enoyl reductase; G – GATase1-like domain. (**B**) Color-coded segmented cryo-EM density map of the core of the FERRY complex, comprising Fy-2 (blue), Fy-4 (red) and Fy-5 (green). The C2-symmetric complex has a clamp-like three-dimensional shape with two arms (Fy-2, Fy-5) extending from a central body (Fy-4). To highlight the different positions of the two Fy-5 molecules in each arm, proximal and distal Fy-5 are colored in dark and light green, respectively. (**C**) Rotated views of the atomic model of the FERRY complex with subunits colored according to (B). (**D**) Rotated views of individual subunits of the FERRY core with Fy-2, Fy-4 and Fy-5 shown on the left, middle and right, respectively. In case of Fy-2 and Fy-4, the dimeric partner is indicated as transparent ribbon representation. The relative location within the complex is highlighted in the cartoon representation below. See also Fig. S1-3 and Table S1, 2.

## Results

### Architecture of the FERRY Rab5 effector complex

In order to understand the interaction of the five subunits of FERRY in molecular detail, we expressed and purified the subunits as described in Schuhmacher et al. (Schuhmacher et al., 2022) (Methods) and reconstituted the complex *in vitro* (Fig. S1). We then determined the cryo-EM structure of FERRY to an overall resolution of 4.0 Å, applying C2 symmetry (Fig. 1B, Fig. S2, 3, Table S1). The structure reveals that the core of FERRY is composed of a dimer of Fy-4, two molecules of Fy-2 and four copies of Fy-5, resulting in a 2:2:4 stoichiometry (Fig. 1B, Fig. S2). The other two subunits, namely Fy-1 and Fy-3, were not resolved in the structure, although SDS-PAGE analysis clearly confirmed their presence in the complex (Fig. S1). The mass of the FERRY complex determined by mass photometry ((Schuhmacher et al., 2022) and Fig. S5) together with the intensities of the corresponding signals of a Coomassie stained SDS-PAGE gel (Schuhmacher et al., 2022) suggests that only a single Fy-1 and Fy-3 bind to the dimeric arrangement of the FERRY core, resulting in a ratio of 1:2:1:2:4 (Fy-1:Fy-2:Fy-3:Fy-4:Fy-5) for the whole complex.

The high quality of the cryo-EM map allowed us to build atomic models for Fy-2 and Fy-4 into the corresponding densities (Fig. 1B-D, Table S1). Densities corresponding to the four Fy-5 molecules, located at the periphery of the reconstruction, exhibited lower local resolution. To obtain an atomic model for these regions, we initially solved the X-ray structure of Fy-5 at 2.7 Å resolution (Table S2) and subsequently relaxed it into the density (Fig. 1C, D, Fig. S3, Table S1, 2).

The structure of the FERRY core reveals an overall clamp-like architecture with two arm-like appendages emanating in opposite directions from a central bulky body (Fig. 1B, C, Fig. S2, 3). Two Fy-4 molecules assemble as a symmetrical dimer, forming the central body of FERRY (Fig. 1D). Each monomer adopts a Rossmann-like fold, where 6 β-strands and 6 α-helices alternate. Through dimerization a continuous 12-stranded twisted ²-sheet encased by two layers of α-helices is formed which is a typical feature of dimeric enoyl reductases. The N-terminal part of Fy-4 is homologous to the catalytic domain of enoyl reductases. Although Fy-4 is equipped with the full set of catalytic residues for enzymatic activity, the location in the center of the complex embraced by helices of Fy-2 blocks the substrate binding site. Thus, a potential catalytic activity of Fy-4 is inhibited in this conformation. However, since human enoyl reductases are normally found as part of the fatty acid synthase complex (Maier et al., 2006) which catalyzes the last step of fatty acid synthesis, a similar enzymatic function of Fy-4 in the context of the FERRY complex seems rather unlikely.

The arm-like appendages of FERRY are formed by a 6-helix bundle of Fy-2 and a Fy-5 dimer each (Fig. 1B-D). The Fy-2 molecules embrace the central Fy-4 dimer and dimerize at their N-and C-terminal ends as coiled-coils, of which only the stem is resolved to high resolution (Fig. 1B-D). The unique architecture of FERRY, which is likely intimately linked to its function as mRNA transport vehicle, does not resemble any of the other large Rab effector complexes, suggesting that FERRY represents a novel class of multi-protein Rab effector complex.

### Fy-2 serves as central scaffolding protein

Fy-2 adopts an integral position within the FERRY complex, as it directly interacts with all other subunits. In the FERRY core, Fy-2 connects Fy-4 and Fy-5 and thereby acts as a scaffold for assembly of the whole protein complex (Fig. 2A, B). The 6-helix bundle of Fy-2 (aa 246-498) is folded in such a way that its antiparallel α-helices form a 5 nm-long hollow tube (Fig. 2A).

**Figure 2.**
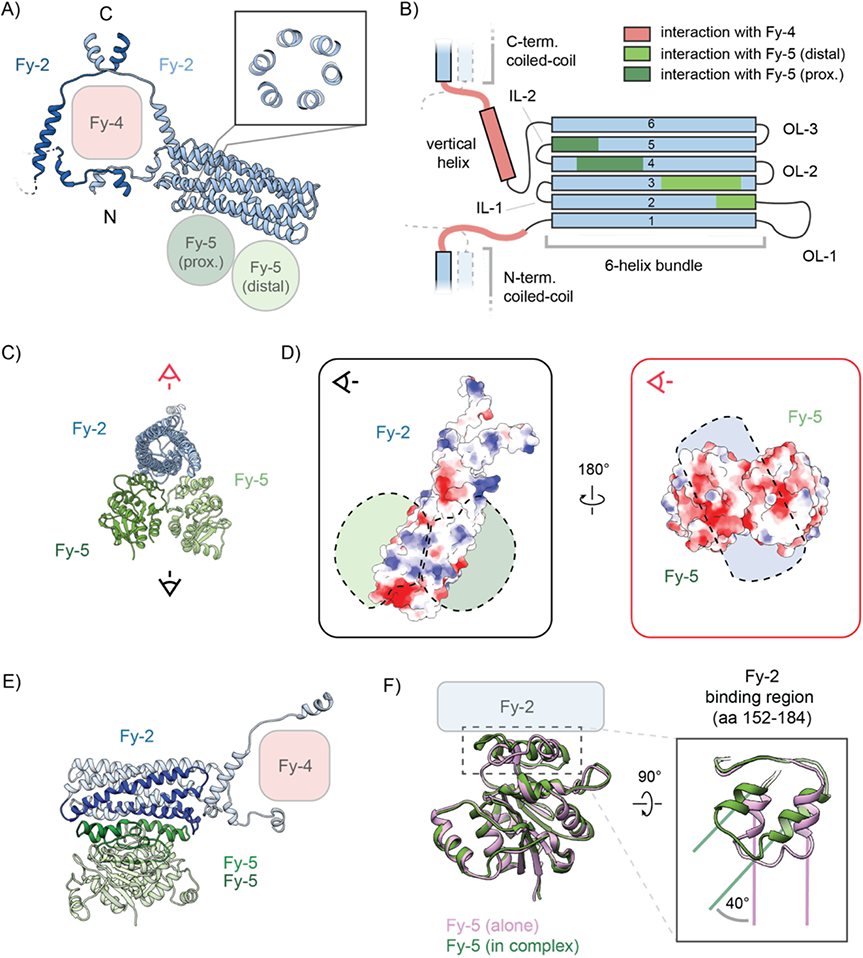
Interaction of Fy-2 with Fy-5. (**A**) Fy-2 adopts an integral position within the FERRY core by interacting not only with the Fy-5 dimer and Fy-4, but it also dimerizes with the second Fy-2 subunit to form two coiled-coil regions. To easier distinguish between the two Fy-2 molecules, they are colored in dark and light blue hues. Notably, the two terminal coiled-coils of Fy-2 extend in opposite directions. The most characteristic feature of Fy-2 is its 6-helix bundle domain, connecting the two coiled coils. Inset shows a cross-section through the 6-helix bundle, highlighting the distinct hexagonal arrangement of the six helices in this region. Same color code as in Fig. 1. (**B**) Topology diagram depicting the domain organization of Fy-2. Interaction regions are highlighted by respective colors. (**C**) The Fy-5 dimer binds to the outer surface of the 6-helix bundle domain of Fy-2. Same color code as in Fig. 1. (**D**) A bottom view, as indicated by the black eye in (C), is depicted on the left. The electrostatic surface of Fy-2 is shown and the position of the Fy-5 dimer indicated as dashed silhouette. In the corresponding 180°-rotated top view (right panel, red eye in (C)), the electrostatic surface of Fy-5 is presented with the position of Fy-2 indicated as dashed silhouette. The predominant positively charged surface of Fy-2 matches the complementary negatively charged surface of Fy-5 in their binding interface. (**E**) Interacting regions of Fy-2 and Fy-5 based on H-D exchange mass spectrometry are highlighted in dark blue and green, respectively. The planar binding surface of Fy-5 exhibited lower deuterium exchange rates when bound to Fy-2, in which the complementary regions of its 6-helix bundle domain are less accessible. (**F**) Superposition of the crystal structure of Fy-5 (pink) and proximal Fy-5 bound in the FERRY complex (green). Inset shows a rotated close-up of the Fy-2-binding region of Fy-5 in which an approx. 40° in-plane rotation is observed upon binding to Fy-2. See also Fig. S4.

The interaction of Fy-2 with Fy-5 is mediated by the 6-helix bundle (Fig. 2B, C). Similar to Fy-4, Fy-5 contains a Rossmann-like fold, composed of a central 6-stranded β-sheet surrounded by six α-helices, and dimerization of Fy-5 results in a continuous 12-stranded twisted β-sheet (Fig. 1D). The proximal and distal Fy-5 subunits bind to helices 4-5 and 2-3 of the 6-helix bundle, respectively, and form a relatively planar interface (Fig. 2B, C). Interestingly, although the 6-helix bundle is non-symmetrical, the interface between the Fy-5 dimer and the 6-helix bundle has a pseudo-two-fold symmetry (Fig. 2C, Fig. S3). The interaction between Fy-5 and the 6-helix bundle of Fy-2 is dominated by charge complementarity (Fig. 2D). While Fy-2 is enriched in positively charged residues at the interface, Fy-5 features a negatively charged patch. These findings are further corroborated by HDX-MS studies showing reduced deuterium exchange in the Fy-2 binding region of Fy-5 as well as in most of the corresponding binding regions of Fy-2 (Fig. 2E, Fig. S4).

To find out whether the interaction of Fy-5 with Fy-2 alters its conformation, we compared our 2.7 Å crystal structure of Fy-5 with Fy-5 in the cryo-EM structure (Fig. 2F, Table S2). Similar to the cryo-EM structure of the complex, Fy-5 formed symmetric dimers and the structure of the major part of the protein was identical (RMSD: 0.812 Å, Fig. 2F). However, the Fy-2 binding region of Fy-5 (aa 152-184) was rotated in-plane by ∼ 40°. Interestingly, this rotation happens in both subunits, so that the dimer stays symmetric and increases the binding interface between the two proteins.

The regions flanking the 6-helix bundle (aa 226–245 and aa 512–540) closely interact with the Fy-4 dimer by wrapping it with extended linkers, including a prominent vertical helix (Fig. 2A, B). Both linkers localize either to clefts or grooves at the protomer-protomer interface of the Fy-4 dimer forming tight interactions based on charge as well as shape complementarity (Fig. 3A, B). This complementarity is particularly prominent in the C-terminal linker where it even extends all the way to the start of the C-terminal coiled-coil domain. A striking feature of the interface between the N-terminal linker and Fy-4 are two electrostatic clusters with up to three different subunits participating. The first cluster comprises Lys-225 and Glu-224 from one Fy-2 molecule, Asp-230 of the second Fy-2 and Lys-299 of Fy-4 (Fig. 3B). The second cluster is formed by Lys-232 of Fy-2 and Asp-306, Glu-309 and Lys-310 of Fy-4.

**Figure 3.**
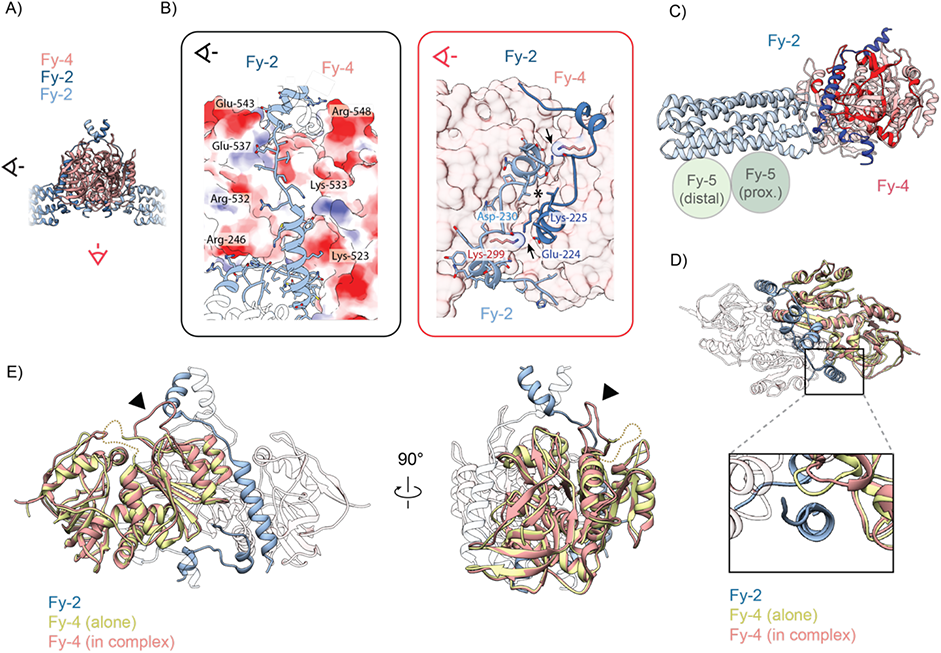
Interaction of Fy-2 with Fy-4. (**A**) Fy-2 interacts with Fy-4 by wrapping around it, before dimerizing with the second Fy-2 to form coiled-coil regions. (**B**) Close-up of the side view, indicated by black eye in (A), is depicted on the left. The Fy-4 dimer is presented as electrostatic surface, interacting parts of Fy-2 as blue ribbon. The vertical helix of Fy-2 is accommodated inside a cleft formed by the two Fy-4 molecules, binding to complementary-charged regions on the surface of Fy-4. A second close-up of the top view, indicated by a red eye in (A), is shown on the right. Fy-4 dimer is shown as semi-transparent surface, Fy-2 as ribbon in light and dark blue with important residues highlighted. The beginning of the N-terminal coiled-coil domain of Fy-4 is highlighted by an asterisk. The complex is stabilized by two charged clusters (black arrows) that are flanking the coiled-coil, containing charged residues from both Fy-2 subunits as well as Fy-4, and by shape complementarity. (**C**) Interacting regions of Fy-2 and Fy-4 based on H-D exchange mass spectrometry are highlighted in dark blue and red, respectively. In Fy-4, mostly regions that constitute the binding cleft for the vertical helix upon Fy-2 binding are not accessible for deuterium exchange. In the case of Fy-2, the vertical helix and regions wrapping around Fy-4 exhibit lower exchange rates. (**D**) Top views of the superposition of the X-ray structure of Fy-4 (yellow) and Fy-4 bound in the FERRY complex (red). Inset shows a close-up of the binding cleft for the vertical helix of Fy-2 (blue), formed by the two Fy-4 protomers. Upon Fy-2 binding, a loop region of Fy-4 moves sideways to accommodate the vertical helix of Fy-2. (**E**) Rotated views of the superposition of the crystal structure of Fy-4 (yellow) and Fy-4 bound in the FERRY complex (red). When Fy-4 is bound in the complex, a previously disordered loop region, indicated as dotted line, becomes ordered (black arrow) and interacts with Fy-2. See also Fig. S4.

The tight interaction between Fy-2 and Fy-4 becomes also evident from HDX-MS measurements (Fig. 3C, Fig. S4). Except for residues 237 to 250, for which no peptides were detected, all parts of Fy-2 that are in close contact with Fy-4 showed a decrease in deuterium uptake in the presence of Fy-4, indicating lower accessibility upon complex formation. The same is true for regions forming the binding clefts of Fy-4.

To more closely investigate the effect of Fy-2 binding to Fy-4, we crystallized the Fy-4 dimer in the absence of Fy-2, solved its structure at 2.9 Å resolution and compared it with our cryo-EM structure of the complex (Fig. 3D, E, Table S2). While the overall structural similarity is high (RMSD: 0.766 Å), we observed two major differences between the two structures. First, the loop of Fy-4 (aa 293-297) moves slightly sideways to accommodate the vertical helix of Fy-2 upon complex formation (Fig. 3D). The second and more striking difference occurs in the region 225-239 of Fy-4, located close to the C-terminal coiled-coil of Fy-2. Here, a previously flexible loop of Fy-4 moves towards Fy-2 to form an additional interface, thereby strengthening the interaction between the two molecules (Fig. 3E).

The N-and C-terminal regions of the two Fy-2 subunits, namely upstream of aa 226 and downstream of aa 540, engage with their respective counterpart to form elongated coiled-coil structures that extend from the complex in diametrically opposite directions. However, the absence of a clear density for the majority of the coiled-coils in our structure indicates flexibility of these domains.

### Role of the two terminal coiled-coils of Fy-2

Our cryo-EM structure revealed the architecture of the FERRY complex core, in which Fy-2 plays a key role as central scaffolding protein. However, two complex subunits, namely Fy-1 and Fy-3, were not resolved in the cryo-EM structure. Data from integrated protein-protein interaction network tools such as STRING indicated a close spatial and functional connection between the two subunits (Szklarczyk et al., 2015). To identify their position within the complex, we performed HDX-MS measurements in the presence and absence of various complex subunits (Fig. 4A, Fig. S4). This allowed us to narrow down the binding regions of Fy-1 and Fy-3 to residues 646-705 of Fy-2, which is located in its C-terminal coiled-coil. We further performed protein cross-linking experiments of the FERRY complex using the zero-length cross-linker 1-ethyl-3-[3-dimethylaminopropyl]carbodiimide hydrochloride (EDC) (Fig. S5A-C). The experiments revealed several positions on Fy-1 and Fy-3 that overlap with the ones determined by HDX-MS. In addition, we observed cross-links from both Fy-1 and Fy-3 to a region adjacent to the main binding site on Fy-2, indicating that Fy-1 and Fy-3 likely form additional contact sites that were not detected by HDX-MS. Furthermore, differential deuterium uptake profiles in the presence and absence of the small GTPase Rab5 allowed us to delineate the Rab5 binding region on the FERRY complex. Like Fy-1/3, it is located on the C-terminal coiled-coil of Fy-2, but a bit closer to the C-terminus (aa 728-752). In known structures of Rab5/Rab5-effector complexes, including those of Rabaptin-5 and EEA1, Rab5 binds via its switch and inter-switch regions either directly to coiled-coils or to regions in their proximity (Mishra et al., 2010; Zhu et al., 2004). Indeed, the Rab5 binding region in the FERRY complex is predicted to form a parallel coiled-coil, further corroborating our HDX data (Fig. 4A, B, Fig. S4). Due to the symmetric structure of the coiled-coil, either one or two Rab5 proteins can bind simultaneously to FERRY.

**Figure 4.**
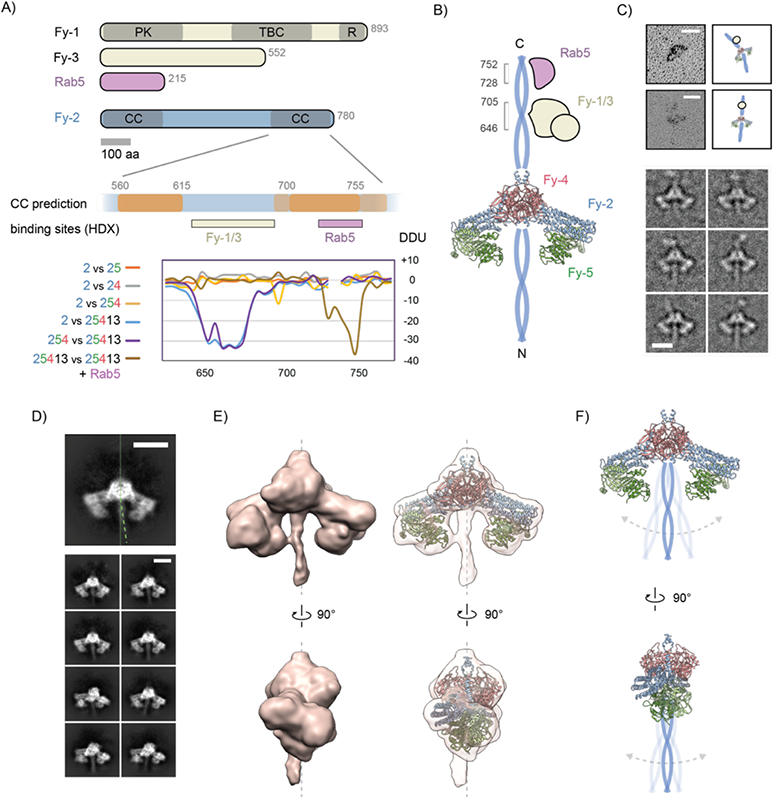
Terminal coiled-coil regions of Fy-2 extend in opposite directions. (**A**) Domain architecture of the Fy-2-interacting proteins Fy-1, Fy-3 and Rab5 is provided at the top. Enlargement of the C-terminal domain of Fy-2 shows predicted coiled-coil regions with orange indicating high CC formation probability. Hydrogen-deuterium exchange mass spectrometry (HDX-MS) measurements delineate the binding regions of Fy-1/3 and Rab5 on the C-terminal coiled-coil of Fy-2. Notably, the Fy-1/3 binding site is in close proximity to the Rab5 binding site, which is located close to the C-terminus of Fy-2. Abbreviations: DDU – differential deuterium uptake; see also schematic in (B). (**B**) Schematic representation of the FERRY complex, displaying that both N- and C-terminal coiled-coil regions of Fy-2 extend in opposite directions from the FERRY core. Binding sites for Fy-1/3 and Rab5 are derived from HDX-MS data. (**C**) FERRY complex visualized by electron microscopy after glycerol spraying and low-angle platinum shadowing is shown on the left with corresponding cartoon illustration provided on the right (upper panel). Two rod-like protrusions extend in opposite directions from the FERRY core. Scale bar: 20 nm. Selected 2D cryo-EM classes of FERRY after hierarchical classification shows density corresponding to the C-terminal coiled-coil region of Fy-2 located at the top of the complex (bottom panel). Scale bar: 10 nm. (**D**) Cryo electron microscopy 2-D class averages of the FERRY complex showing that the N-terminal coiled-coil of Fy-2 can adopt multiple position relative to the FERRY core. Scale bar: 10 nm. (**E**) Rotated views of 3-D reconstruction of FERRY without applied symmetry, filtered to 15 Å. Right panel shows fitting of the atomic models for the FERRY core into the density for orientation. The elongated density that protrudes from the central Fy-4 dimer corresponds to the N-terminal coiled-coil of Fy-2. The deviation from the central complex axis suggests a certain degree of flexibility for the N-terminal coiled-coil. (**F**) Schematic representation of the FERRY complex highlighting the flexible nature of the N-terminal coiled-coil of Fy-2. Both “arms” (6-helix bundle domain of Fy-2 and the Fy-5 dimer) appear to restrict the degree of movement.See also Fig. S4, 6 and Movie S1, 2.

In our cryo-EM structure we could only identify density corresponding to the first few residues of the C-terminal coiled-coil region of Fy-2 (Fig. 1B, Fig. S2, 3). This suggests a high flexibility of the rest of the coiled-coil, which is a common observation also for other coiled-coil-containing complexes (Shi et al., 2020). In general, long coiled-coils without a rigidifying interaction partner are mostly flexible and thus challenging to visualize in EM. To visualize the C-terminal coiled-coil region as well as the Fy-1 and Fy-3 subunits, we performed low-angle platinum shadowing experiments with the FERRY complex and extended hierarchical clustering of selected initial cryo-EM FERRY classes (Fig. 4C, Fig. S6, Movie S1). In the former analysis, we observed particles with two rod-like protrusions emanating from opposite sides of the FERRY core (Fig. 4C, Fig. S6). These protrusions probably correspond to the two terminal coiled-coil regions of Fy-2. The extended classification of the cryo-EM data revealed a rod-like density and a globular density that can be attributed to the coiled-coil and possibly to the Fy-1/3 subunits, respectively (Fig. 4C, Fig. S6, Movie S1).

In the case of the N-terminal coiled-coil of Fy-2, we observed only density corresponding to the beginning of the coiled-coil at high-resolution (Fig. 1B, Fig. S2, 3), similar to its C-terminal counterpart. However, at lower thresholds and in particular in 2D class averages we could also identify density in the space between the two arm-like appendages (Fig. 4D). This density can be unambiguously identified as a coiled-coil structure which adopts multiple orientations relative to the FERRY core (Fig. 4D, Movie S2). This also explains the lower resolution in this region of the complex (Fig. S3). Therefore, we calculated a reconstruction of the FERRY complex without applying symmetry, yielding a 6.2 Å map (Fig. 4E, Fig. S2, 3). Although the resolution of the coiled-coil did not improve and still appeared only at a lower threshold, its rod-like appearance including a twist is reminiscent of a coiled-coil. We believe that the proximal Fy-5 molecules bound to the arm restrict the overall mobility of the N-terminal coiled-coil of Fy-2 (Fig. 4F). The functional implications of this restricted degree of movement, however, are not yet understood. Taken together, our results show that both terminal coiled-coil domains of Fy-2 have different degrees of flexibility and act as a binding hub for other FERRY subunits. This even further underpins the central role of Fy-2 in the FERRY complex, as it directly interacts with all other four members of the complex as well as with Rab5.

### Interaction of FERRY with mRNA

To determine whether the purified FERRY complex binds mRNA *in vitro*, we performed electrophoretic motility shift assays (EMSA) with the fully reconstituted FERRY complex (Fy-1/2/3/4/5) and four different mRNAs (*mrpl41, mdh2, prdx5, pigl*) (Fig. 5B, Fig. S7A). The results clearly show that all different mRNAs bind to FERRY. Interestingly, with increasing amounts of mRNA the signal corresponding to the FERRY-RNA complex sharpens into a band and migrates at a lower molecular weight (Fig. 5B-D, Fig. S7A-C). This is an indication that RNA binding decreases the structural flexibility of the complex.

**Figure 5.**
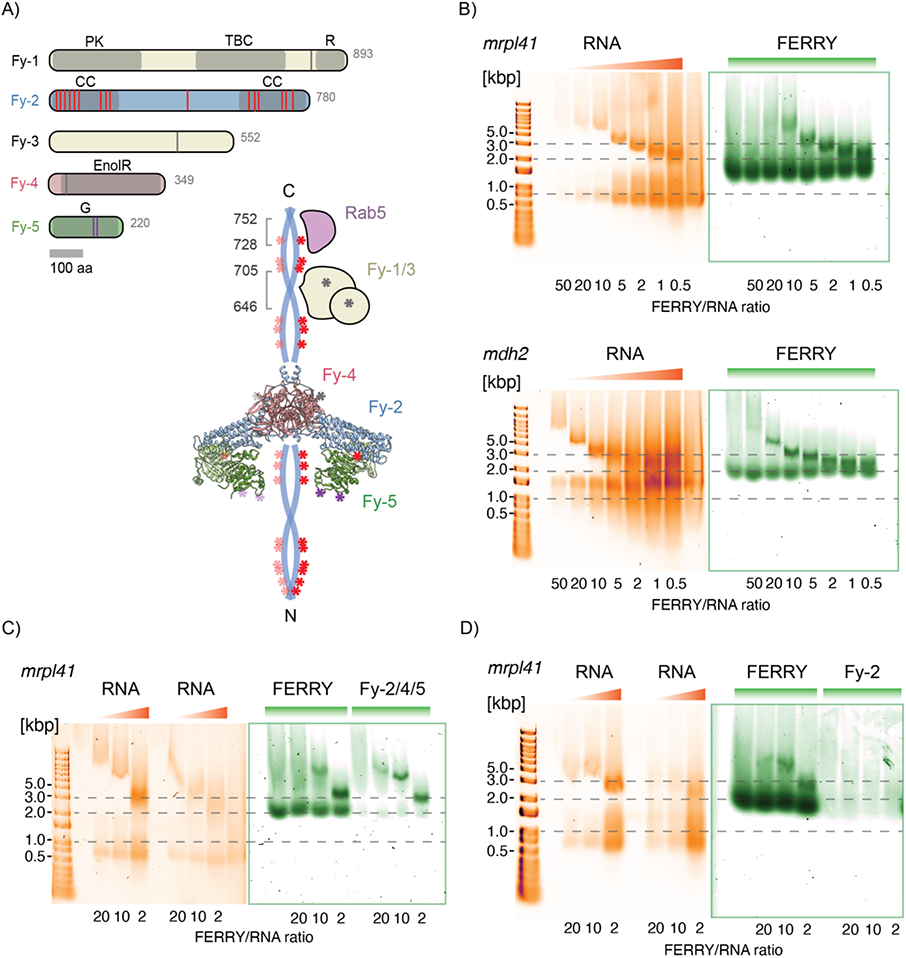
RNA binding of the FERRY complex. (**A**) In the lower right panel, positions of UV cross-links between RNA/Fy-2 and RNA/Fy-5 are indicated in the FERRY complex by red and lilac asterisks, respectively. Crosslinks with RNA in the other FERRY subunits are indicated by grey asterisks. Only crosslinks are shown that were identified in at least 3 out of 5 replicates. Notably, crosslinks are enriched at the terminal coiled-coils of Fy-2. Together with Fy-5, the N-terminal coiled-coil of Fy-2 constitutes a clamp-like structure, which is important in RNA binding. In the upper left panel, the domain organization of FERRY subunits is depicted with positions of crosslinks indicated. Domain abbreviations: PK – Pseudokinase; TBC – Tre-2/Bub2/Cdc16; R – Rhodanese; CC – Coiled coil; EnolR – Enoyl reductase; G – GATase1-like domain. (**B**) Electrophoretic mobility shift assays (EMSAs) showing the interaction between the FERRY complex and *mrpl41* and *mdh2* mRNA in the upper and lower panel, respectively. RNA is stained with SYBR Gold (orange, left) and protein with Sypro Red (green, right). Note that with increasing ratios of RNA:FERRY a shift towards higher mobility of the FERRY-RNA complex is observed. (**C**) EMSAs showing that both, FERRY and the 3-subunit Fy-2/4/5 complex, are capable of binding the mRNA *mrpl41 in vitro*. (**D**) EMSAs showing that in contrast to the fully assembled FERRY complex, Fy-2 alone is not sufficient *mrpl41* mRNA. See also Fig. S7 and File S1.

A protein structure comparison using the DALI server found several members of the DJ-1/ThiJ/PfpI superfamily to have a similar structure to Fy-5. Interestingly, DJ-1, which shares the Rossmann-like fold and overall structure with Fy-5 (RMSD: 1.012 Å, Fig. S3), has been shown to bind RNA at nanomolar concentrations, and mutations in the corresponding gene have been linked to neuronal degeneration (Brug et al., 2008; Lee et al., 2003). The close structural similarity between both proteins suggests that Fy-5 could perform a similar function, *i.e.* contribute to RNA binding, in the FERRY complex. However, EMSAs showed that Fy-5 alone does not exhibit detectable RNA binding ability (Fig. S7D). We can envisage two possible explanations: Either Fy-5 is not involved in mRNA interaction, or the interaction interface is more complex and requires different subunits of the FERRY complex simultaneously. In order to resolve this question, we performed UV-induced protein-RNA crosslinking mass spectrometry (Kramer et al., 2014) (Fig. 5A, SI File 1). The mass spectrometry analysis revealed in total 37 lysine residues within Fy-1 to Fy-5 crosslinked to RNA (SI File 1). Only two of the crosslinked lysine residues are found in one loop region of Fy-5 (Fig. 5A). Unexpectedly, most crosslinks cluster along the coiled-coils of Fy-2 instead. This suggests that bound RNA stretches over the whole length of the FERRY complex or that several RNA molecules bind simultaneously to the same FERRY complex at different positions. Interestingly, several crosslink sites are located in the cavity of the clamp-like structure in the FERRY complex, suggesting that this clamp plays an important role in mRNA binding (Fig. 5A). Consistent with the crosslinking data, the bottom of the cavity is lined by positively charged residues which are known to be essential for the interaction of proteins with RNA (Fig. S3) (Ghaemi et al., 2017; Lunde et al., 2007).

In order to verify that Fy-2 is sufficient for RNA binding, we carried out EMSAs with the three-subunit Fy-2, Fy-4 and Fy-5 core complex. RNA bound to this complex as efficiently as to the whole FERRY complex, indicating that Fy-1 and Fy-3 are not required for RNA binding (Fig. 5C, Fig. S7B). However, when we performed the measurements with Fy-2 alone, unlike in the RNA cross-linking studies, we observed that RNA binding was almost entirely abrogated (Fig. 5D, Fig. S7B). It is possible that Fy-2 alone may not be able to adopt the correct conformation/folding to fulfil its function as RNA interactor as in the presence of Fy-4 and Fy-5.

To test this hypothesis and determine whether the coiled-coil domains of Fy-2 constitute indeed the main RNA binding site, we reconstituted different FERRY complexes of Fy-2 variants, in which the N-terminal coiled-coil had been gradually truncated (Fig. 6A, B, D). EMSAs of these FERRY complexes showed that their RNA binding ability decreased almost proportionally with the length of the coiled-coil region, indicating that the coiled-coils indeed comprise the main RNA binding site of Fy-2 (Fig. 6F). Thus, Fy-4 and Fy-5, which do not exhibit detectable RNA binding ability on their own (Fig. S7D), likely act as a stabilizer and scaffold for the proper orientation of the coiled-coils of Fy-2.

**Figure 6.**
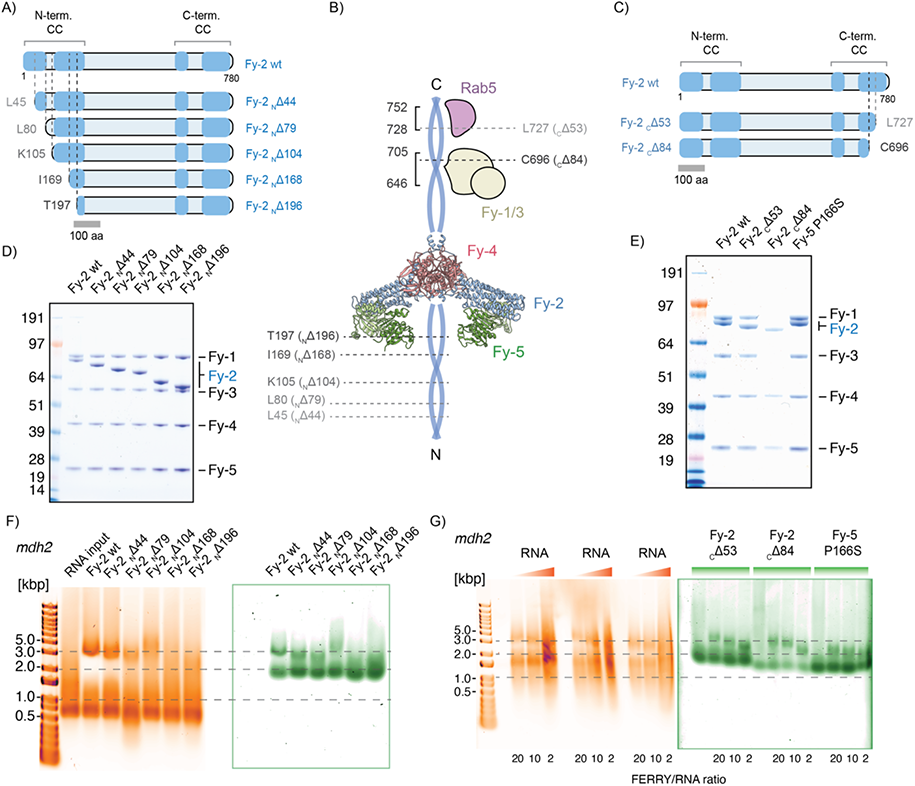
Terminal coiled-coil domains of Fy-2 are essential for RNA binding. (**A**) Overview of N-terminal truncation variants of Fy-2. Predicted coiled-coil regions are highlighted in blue. (**B**) Schematic representation of the FERRY complex with positions of N-and C-terminal truncations indicated by dashed lines. (**C**) Overview of C-terminal truncation variants of Fy-2. Predicted coiled-coil regions are highlighted in blue. (**D-E**) SDS-PAGE of reconstituted FERRY complex having the wt Fy-2 subunit replaced by either N-terminal (D) or C-terminal (E) truncation variants. (**F-G**) Electrophoretic mobility shift assays (EMSAs) showing the interaction between *mdh2* mRNA and the FERRY complex with Fy-2 being replaced by either N-terminal (F) or C-terminal (G) truncation variants. RNA is stained with SYBR Gold (orange, left) and protein with Sypro Red (green, right). With decreasing length of the N-terminal coiled coil, Fy-2 loses its ability to bind RNA. The two C-terminal truncation variants as well as the P166S point mutation do not affect RNA binding significantly. See also Fig. S5, 7.

Together, these results demonstrate that although Fy-5 is part of a complex mRNA binding interface on the FERRY complex, the majority of interactions are mediated by the coiled-coils of Fy-2, which are structurally stabilized by Fy-4 and Fy-5.

To assess the number of mRNA molecules bound to FERRY, we performed mass photometry experiments with FERRY in the absence and presence of *mdh2* RNA (Fig. S5E). We measured an overall mass of 501 +/- 25 kDa for the FERRY complex alone, corresponding well to the calculated molecular weight of 521 kDa. Considering that the mass was estimated based on protein standards, the measured molecular weight of mdh2 RNA alone is as well close to the calculated value (722 kDa vs. 655 +/- 29 kDa). Upon FERRY-RNA complex formation, we observed an additional peak at 1282 +/- 52 kDa, likely corresponding to a 1:1 FERRY-mdh2 RNA complex. Since we did not observe peaks at higher molecular mass, we conclude that *in vitro* one RNA molecule binds to a FERRY complex at a time.

### Disease-relevant mutations of FERRY

Our structural and functional knowledge of FERRY allows us to analyze further the molecular basis of disease-associated mutations of the FERRY complex. In a 2018 study, the homozygous nonsense variant c.2089C>T (p.Arg697*), which results in a truncated Fy-2 protein lacking the last 84 residues was identified in a patient with a developmental and neurological disorder (Suleiman et al., 2018). To investigate whether this mutation affects Fy-1/3 binding and the interaction with Rab5 as suggested by our HDX-MS studies (Fig. 4A), we reconstituted and analyzed two FERRY variants with C-terminal deletion mutations in Fy-2, *i.e.* Fy-2 _C_Δ53 and Fy-2 _C_Δ84 (Fig. 6B, C, E, G). The first mutant lacks the region that was identified as the Rab5 binding site, whereas the second mutant also partially lacks the region identified as main Fy-1/3 interface (Fig. 6B, C). The reconstitution of both complexes revealed that while Fy-2 _C_Δ53 together with the other proteins formed a stable five subunit complex, Fy-2 _C_Δ84 lacks the ability to interact with Fy-1 and Fy-3, resulting in a three-subunit core complex, i.e. Fy-2 _C_Δ84/Fy-4/5 (Fig. 6E). Both complexes showed no impairment in RNA binding (Fig. 6G).

We additionally performed GST-Rab5 pulldown assays to examine Rab5 binding with the described FERRY mutant complexes (Fig. S5D). While the variant lacking the C-terminal 84 residues completely abrogated Rab5 binding, the Fy-2 _C_Δ53 mutant displayed residual Rab5 binding, indicating that the _C_Δ53 truncation included most but the entire Rab5 interface. This suggests that the Rab5 binding interface identified by HDX (725-752) is indeed correct but must probably be extended by several residues towards the N-terminus. Taken together, the disease-related C-terminal truncation of Fy-2 (_C_Δ84) is impaired in FERRY complex assembly and also unable to bind Rab5, most likely giving rise to severe mis-regulation of mRNA localization and transport.

For Fy-5, the point mutation P166S has been associated with gastric adenocarcinoma (https://cancer.sanger.ac.uk). This particular mutation is located in the Fy-2 binding region of Fy-5 and could hence disrupt complex formation. To better understand the molecular effect of this mutation, we introduced the point mutation (P166S) into Fy-5 and reconstituted a FERRY complex. We did not observe any impairment of FERRY assembly and RNA binding compared to the wildtype complex *in vitro* (Fig. 6E, G, Fig. S7C). This could mean that the disturbances caused by this mutation are too subtle to be picked up by our *in vitro* assays or that they interfere with other processes unrelated to complex formation and mRNA transport *in vivo*. The association to gastric adenocarcinoma would argue as such, since aberrant mRNA localization or transport mainly causes neuronal symptoms and brain malfunction.

### Hydrophobic cavity in the 6-helix bundle of Fy-2

The 6-helix bundle of Fy-2 is a rather rare structural arrangement, connecting Fy-4 with the Fy-5 dimer (Fig. 7A). To our surprise, we identified an elongated density within the 6-helix bundle, located at its Fy-4-facing side (Fig. 7B). While the exterior of the 6-helix bundle is mostly polar, its interior is highly enriched with hydrophobic residues (Fig. 7B, C). This suggests that the elongated density, which is centered almost perfectly within the hydrophobic cavity, *i.e.* residing on the central tube axis, probably corresponds to a hydrophobic molecule. When we analyzed the arrangement of the six helices that constitute the tube in more detail, we observed a change in their organization along the central axis (Fig. 7D). Starting from a hexagonal pattern on the Fy-4-facing side, they transition en route towards a more pyramidal-like arrangement on the opposite end. This rearrangement is accompanied by a gradual decrease of the intraluminal diameter of the tube. Consequently, the putative molecule could not access the tube from the ‘pyramidal’ side unless major rearrangement occurred in the 6-helix bundle. To enter from the Fy-4-facing side therefore appears to be the more likely scenario, but would still require the displacement of one or more nearby loop regions of the 6-helix bundle, i.e. IL-1 and/or IL-2, in order to grant access to the hydrophobic interior. Another possible function of the hydrophobic moiety could be to support and facilitate the assembly of the FERRY complex, for example by stabilizing the 6-helix bundle of Fy-2.

**Figure 7.**
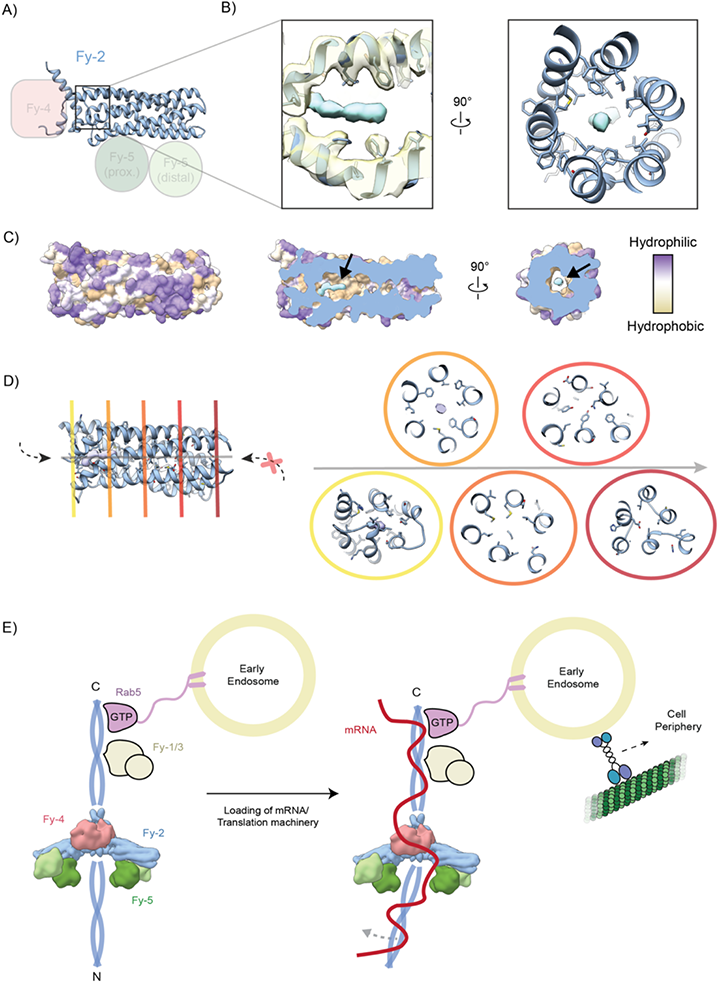
Hydrophobic binding pocket and Model of FERRY recruitment and RNA binding. (**A**) The 6-helix bundle domain of Fy-2 adopts a crucial position within the FERRY complex. (**B**) The left inset shows an enlarged side view of the interior of the 6-helix bundle domain at the Fy-4-facing end of the tube, as indicated in (A). Both, 3D-reconstruction (semi-transparent) and fitted atomic model (blue) are shown. An additional, elongated density (cyan) is accommodated within the hollow cavity formed by the 6-helix bundle. A rotated view shown in the right panel further highlights the almost perfectly centered position of the cyan density as well as the hydrophobic environment that is created by inward-facing hydrophobic side chains. (**C**) Surface hydrophobicity of the 6-helix bundle domain of Fy-2 is depicted on the left, showing that the cytoplasm-exposed exterior is primarily hydrophilic. Cross sections of different orientations in the middle and right panel highlight the hydrophobic nature of the interior cavity, with the position of the elongated density indicated by an arrow. (**D**) Different color-coded cross sections of the 6-helix bundle domain of Fy-2 are shown in the right panel with their respective position along the central tube axis (grey) indicated in the left panel. The six helices transition from a more ordered hexagonal arrangement (yellow, closest to Fy-4) to a more pyramidal-like array (red, opposite end). This en route transition correlates with a decrease of the interior space, causing a narrowing of the cavity towards the distal end. (**E**) Model of FERRY recruitment and RNA binding: The activated GTP-bound form of Rab5 recruits the FERRY complex by binding to the C-terminal coiled-coil domain of Fy-2. Since Rab5 proteins insert via two C-terminal lipidated cysteines into the early endosome (EE) membrane, FERRY also becomes associated with the EE. FERRY interacts with mRNA and/or the translation machinery, with a single RNA molecule bound to the coiled-coils of Fy-2. The flexibility of the N-terminal coiled-coil might facilitate RNA binding via the FERRY clamp. Ultimately, motor protein-mediated transport of EEs via the MT network delivers the FERRY-associated RNA cargo to its respective cellular target destination.

The implications of this unidentified molecule on the function of the FERRY complex are not immediately obvious and represent a compelling topic for further investigations.

## Discussion

In this study, we resolved the core of the FERRY complex to 4.0 Å resolution using single particle cryo-EM. The structure is composed of a central Fy-4 dimer of which two arms, each consisting of a Fy-2 molecule and a Fy-5 dimer, protrude in different directions. In the scaffold protein Fy-2, both terminal regions interact with their symmetry-related counterpart to form flexible coiled-coils that extend in opposite directions. Furthermore, HDX measurements, cross-linking mass spectrometry, mutational studies and GST-pulldowns allowed us to delineate the binding sites of the subunits Fy-1 and Fy-3 as well as Rab5, all of which are located on the C-terminal coiled-coil of Fy-2.

The cryo-EM structure in combination with corresponding HDX- and cross-linking mass spectrometry provides important insights into the architecture of the FERRY complex and the interaction between its five subunits. Although the structural and functional record is by no means complete for the long-distance RNA transport via EEs, we can use the information provided by our atomic model of the FERRY complex to define critical steps in the process and suggest the following mode of FERRY recruitment and loading (Fig. 7E). Activated EE-associated Rab5 recruits the FERRY complex through binding to the C-terminal coiled-coil region of Fy-2 to the EE. The Rab5 binding site locates adjacent to the Fy-1 and Fy-3 subunits, which also bind to the C-terminal coiled-coil region. Since there is no obvious additional binding site for EE proteins on FERRY, we believe that the elongated complex sits with its long axis at 90° to the EE surface, although it should be borne in mind that the connection of the Rab5 globular domain to the membrane is not likely to be rigid due to the poorly ordered long hypervariable domain of the GTPase C-terminus in the absence of further interactions. Mutations in Fy-1 cause intellectual disability and severe infantile syndromic encephalopathy in patients, both characterized by brain atrophy, highlighting its importance in FERRY-mediated RNA-transport (Bhoj et al., 2016; Chong et al., 2016). In general, perturbations in FERRY-mediated long-range RNA transport, which is particularly relevant to neurons, typically manifest in brain disorders. As for Fy-1, mutations in Fy-3 have been linked to intellectual disability, based on genetic analysis of two Finnish and one Dutch family (Philips et al., 2017).

For Fy-2, several mutations connected to malfunction of the brain have been described including four biallelic loss of function variants that have been linked to neurodevelopmental syndrome (Rehman et al., 2019). Our study allowed us to show that both complex assembly and Rab5 binding are impaired in a homozygous Fy-2 nonsense variant which was identified in a patient with developmental delay and brain abnormalities (Suleiman et al., 2018). This variant lacks the 84 C-terminal residues on Fy-2 which contain the Rab5 as well as Fy-1/Fy-3 binding sites, likely resulting in a failure of FERRY to bind to the EE via Rab5 *in vivo*, further underlining the importance of these subunits for the proper action of FERRY-mediated RNA transport.

For the point mutation P166S in Fy-5 which has been linked to gastric adenocarcinoma, we did not observe any impairment of FERRY assembly and RNA binding compared to the wildtype complex *in vitro*. This indicates that the association with cancer might be connected to functions involving the FERRY complex *in vivo*.

So far, no mutations have been described for Fy-4. There is the possibility that point mutations in this protein, despite its central position in the FERRY complex have less severe effects on the function of FERRY since it serves mainly as scaffold.

Our crosslinking mass spectrometry measurements and EMSA studies demonstrated that RNA binds primarily to the coiled-coils of Fy-2, as well as a loop of Fy-5, indicating that it stretches over the entire length of the FERRY complex. The composite binding interface, comprising different FERRY subunits, might be necessary to create different binding specificities for various mRNA transcripts *in vivo*. The three-dimensional arrangement of possible RNA-binding sites likely determines the specificity and binding affinity of certain RNA molecules. Accordingly, electrophoretic motility shift assays showed that the binding affinities to FERRY varied between different RNA transcripts.

The structure of the FERRY core resembles that of an RNA-binding clamp, which raises the question whether this structure has mechanical implications. We observed a certain level of flexibility in the arms of the clamp in cryo-EM (Fig. S3, Movies S1, 2) indicating that the structural movements necessary for slight opening and closing of the clamp are possible. In addition, the N-terminal coiled-coil of Fy-2 is to a certain degree flexible which could facilitate the binding of the RNA by providing space inside the clamp. However, we propose that rather than mechanically clamping the RNA, the adaptable large three-dimensional cavity of the clamp provides the optimal binding site for structurally flexible RNAs. As shown by our truncation studies, the quite long coiled-coils of Fy-2 provide additional binding sites for RNA, thereby increasing the specificity and/or affinity for the cargo. FERRY binds a single RNA molecule as indicated by our mass photometry data (Fig. 7E). Although we demonstrated that RNA binds directly to FERRY, we cannot exclude that additional adaptor proteins are involved in this process *in vivo*. The long N-terminal coiled-coil of Fy-2 would provide enough space for their binding.

Another question that arises is the function of the density that we identified in the hydrophobic pocket formed by the 6-helix bundle of Fy-2. Due to the hydrophobic environment, we assume that the density corresponds to a hydrophobic molecule. However, based on the shape of the density and its unknown occupational state in the structure, we cannot distinguish whether it is a lipid or single fatty acid/aliphatic chain. We can exclude that it is detergent or a similar amphipathic compound since these were not added during purification. It is also very unlikely that it is a post-translational modification. First of all, there would be no space for a modifying enzyme. In addition, Leu-333, Thr-335 and Val-332 which are in close proximity to the ends of the density are non-typical residues for lipidic modifications.

Binding of hydrophobic molecules to other RNA-binding proteins has been described before, mainly resulting in a modulation of their respective RNA binding activity. Among these molecules is the natural product hippuristanol, first identified in the coral *Isis hippuris*, that inhibits eukaryotic translation (Bordeleau et al., 2006). Potent binding of the steroid locks the eukaryotic elongation factor eIF4A in an aberrant closed conformation, thus preventing it from interacting with RNA (Cencic and Pelletier, 2016). Another example in which RNA binding is impaired by hydrophobic moieties is the stem cell translation regulator Musashi-1. Here, 18-22 carbon ω-9 mono-unsaturated fatty acids can bind to the N-terminal RNA recognition motif, resulting in a conformational change that prevents RNA association (Clingman et al., 2014). We cannot exclude that the unknown hydrophobic molecule exerts a similar inhibitory effect on the RNA binding activity of FERRY. However, this scenario is rather unlikely since the main RNA binding regions of the FERRY complex are located on the terminal coiled-coils of Fy-2 and, therefore, far away from the hydrophobic moiety. In addition, we observed RNA binding to FERRY in the presence of the hydrophobic moiety.

Although highly speculative, it is conceivable that the hydrophobic cavity might accommodate the lipidic tail of prenylated Rab5 and could thus act similar to other GDIs, *i.e.* facilitating the transport of Rab5 through the cytosol. In this way, another layer of regulation of Rab5 activity could be theoretically achieved by the FERRY complex. Since we have not co-purified Rab5 with the FERRY complex, we assume that in our case the binding pocket is occupied by a hydrophobic molecule that was inserted during the recombinant expression.

Taken together, our structural and functional analyses of FERRY extend the understanding of this remarkable Rab5 effector complex and hence also shed light onto how intracellular mRNA transport processes are coordinated within the cell, serving as a basis for future research.

## Material and Methods

### Molecular cloning

The human proteins Fy-1 (Tbck, ENSG00000145348, Q8TEA7), Fy-2 (Ppp1r21, ENSG00000162869, Q6ZMI0), Fy-3 (C12orf4, ENSG00000047621, Q9NQ89), Fy-4 (Cryzl1, ENSG00000205758, O95825), Fy-5 (Gatd1, ENSG00000177225, Q8NB37) and Rab5a (ENSG00000144566, P20339) were used in vectors as described (Schuhmacher et al., 2022). Fy-4 and Fy-5 were expressed with a non-cleavable N-terminal hexa-histidine (His_6_) tag in bacteria and insect cells, respectively. Fy-1, Fy-2 and Fy-3 were combined in multi-gene plasmid with Fy-1 carrying a N-terminal His_6_ tag and produced in insect cells. Rab5 was used as GST fusion variant in a pGEX-6P-3 or pGAT2 vector.

### Virus production and insect cell expression

SF9 cells growing in ESF921 media (Expression Systems) were co-transfected with linearized viral genome and the expression plasmid and selected for high infectivity. P1 and P2 viruses were generated according to the manufacturer’s protocol. Best viruses were used to infect SF9 cells at 10^6^ cells/mL at 1% vol/vol and routinely harvested after 40-48 hours at about 1.5×10^6^ cells/ml. The pellet was suspended in lysis buffer (20 mM HEPES, pH 8.0, 250 mM NaCl, 20 mM KCl, 20 mM MgCl_2_ and 40 mM imidazole) or SEC buffer (20 mM HEPES, pH 7.5, 250 mM NaCl, 20 mM KCl and 20 mM MgCl_2_) supplemented with a protease inhibitor cocktail, flash frozen in liquid nitrogen and stored at −80 °C.

### Protein expression and purification

The FERRY complex, its individual components Fy-4 and Fy-5 and Rab5 were essentially expressed and purified as described in (Schuhmacher et al., 2022). For better readability a brief description of the expression and purification is given in the following.

In general, purified proteins were analyzed using SDS-PAGE and their concentration determined by spectrophotometer (NanoDrop Lite, Thermo Scientific) unless stated.

Fy-5 and GST-Rab5: Fy-5, Fy-5 P166S and GST-Rab5 (from pGAT2) were expessed in *E. coli* BL21 (DE3) under autoinduction conditions using D-(+)-lactose monohydrate at 1.75 % (w/v), supplemented with the respective antibiotics (50 μg/mL kanamycin or 100 μg/mL ampicillin) at 30 °C. Harvested bacteria were suspended in lysis buffer and subsequently lysed or stored at −80 °C. After lysis (sonication) the protein was purified from the clarified lysate in a two-step purification, involving Ni-NTA affinity chromatography (HisTrap FF column, GE Healthcare) and size exclusion chromatography (SEC) (HiLoad 16/600 Superdex 200 pg, GE Healthcare) in SEC buffer.

Fy-4: After sonication and clarification of the lysate by centrifugation (22 500 rpm/61 236 x g, 20 min, 4 °C), the lysate was filtrated using Millex® HV membrane filter units with a pore size of 0.45 µm (Merck Millipore). Fy-4 was subsequently purified combining Ni-NTA affinity chromatography (HisTrap FF column, GE Healthcare) and SEC (HiLoad 16/600 Superdex 200 pg, GE Healthcare).

FERRY complex: To reconstitute the FERRY complex or the respective FERRY variants, Fy-1 to Fy-3 were expressed from a single virus and the harvested insect cells supplemented with purified Fy-4 and Fy-5 prior to cell lysis (Microfluidizer LM20, Microfluidics). The purification was accomplished by combining affinity chromatography and SEC. After clarification (22 500 rpm/61 236 x g, 20 min, 4 °C) and filtration (Millex® HV membrane filter units), the lysate was supplemented with Ni-NTA agarose (Qiagen, 1.3 ml resin/ 1 l insect cell pellet). Subsequently, the resin was transferred into gravity flow chromatography columns (Poly-Prep® Chromatography Column, Bio-Rad) and extensively washed with lysis buffer and wash buffer (20mM HEPES, pH 7.5, 250mM NaCl, 20mM KCl, 20mM MgCl_2_ and 80mM imidazole). The complex was eluted with elution buffer (20 mM HEPES, pH 7.5, 250 mM NaCl, 20 mM KCl, 20 mM MgCl_2_ and 500 mM imidazole) in 1 ml fractions and protein containing fractions were applied to SEC without further concentration using a Superose 6 increase (Superose 6 Increase 10/300 GL, GE Healthcare) which was equilibrated in SEC buffer.

Rab5: Expression of GST-Rab5 was performed under autoinduction conditions as described for Fy-5. Harvested bacterial pellets were resuspended in SEC buffer, lysed using sonication and the lysate clarified by centrifugation (22 500 rpm/61 236 x g, 20 min, 4 °C). GST-Rab5 was captured on Glutathione Sepharose 4B resin (Cytiva), extensively washed with SEC buffer and cleaved off the resin using HRV 3C protease (produced in house). The protein was subsequently concentrated using Amicon Ultracel-30K (Millipore) centrifuge filters and applied to SEC using a Superdex 200 column (HiLoad 16/600 Superdex 200 pg, GE Healthcare) equilibrated in SEC buffer.

Rab5 was loaded with GTPγS prior to the HDX experiments. To do so, Rab5 was concentrated using an Amicon Ultracel-30K (Millipore) centrifuge filter, subsequently supplemented with 2.5 mM GTPγS and 250 nM of a GST fusion of the Rab5 GEF domain of Rabex5 and incubated for 60 mins on ice. To remove the Rab5 GEF domain, Glutathione Sepharose 4B was added to the mixture and incubated for 90 mins at 4 °C. The resin was pelleted by centrifugation (12 000 rpm/ 15 300 x g, 10 min, 4 °C) and the supernatant containing the GTPγS loaded Rab5 was flash frozen and stored at −80 °C. The protein concentration was determined using a BCA assay (Pierce™ BCA Protein Assay Kit, Thermo Scientific).

### GST-Rab5 pulldown assays

Direct GST-Rab5 interaction assays were performed as described in (Schuhmacher et al., 2022). In brief, 12 µL Glutathione Sepharose 4B (Cytiva) was saturated with 5 nmol GST-Rab5 and unbound protein removed. The beads were incubated with 1 mM nucleotide (GDP or GTPγS) and 420 nM of GST-Rabex5-Vps9 for 60 mins at 4 °C to achieve nucleotide loading. After washing, 0.2 nmol of FERRY or the different FERRY variants were added to the beads in a total volume of 100 μl and incubated for 20 min at 4 °C on a shaker (700 rpm). After extensive washing, proteins were eluted with SEC buffer supplemented with 20 mM GSH and analyzed by SDS-PAGE.

### mRNA production and electrophoretic motility shift assays (EMSA)

mRNA sequences for *mdh2*, *mrpl41*, *prdx5* and *pigl* and *in vitro* transcription information are detailed in (Schuhmacher et al., 2022). The mRNAs were produced and purified by *in vitro* transcription using the T7 RiboMAX™ Express Large Scale RNA Production System (Promega) according to the manufacturer’s protocol and subsequent phenol:chloroform extraction and isopropanol precipitation. RNA concentrations were determined by spectrophotometer (NanoDrop Lite, Thermo Scientific) and the RNA was stored at −80 °C until usage.

For direct protein-RNA interaction assays, respective amounts of FERRY complex were mixed with *in vitro* transcribed mRNA in varying protein/RNA ratios in SEC buffer in a total volume of 20 or 35 μl and incubated for 80 min at 37 °C. The samples were analyzed using gel electrophoresis, RNA was visualized using ethidium bromide or SYBR™ Gold Nucleic Acid Gel Stain (Invitrogen) and proteins were stained using SYPRO Red Protein Gel Stain (Sigma Aldrich) according to the manufacturers’ protocols.

### Mass photometry

For the mass photometry measurements 10 pmol FERRY complex was mixed with 10 pmol *in vitro* transcribed *mdh2* RNA in a total volume 20 ml and incubated for 80 mins at 37 °C. The sample was diluted into SEC buffer to reach 100 nM prior to data acquisition. Mass Photometry (MP, iSCAMS) was performed on a TwoMP instrument (Refeyn, Oxford, UK) at room temperature. High precision 24 x 50 mm coverslips (Thorlabs CG15KH) were cleaned with ultrasound, rinsed with isopropanol and water and dried with clean nitrogen gas (Young and Lin, 2018) and coated with 0.01 % Poly-L-lysine solution (Sigma, P4832). Mass calibration was performed using BSA, IgG, Thyroglobulin standards (Sigma), Low DNA Mass ladder (Invitrogen, 10068013) and Millenium RNA marker (ThermoFisher, AM7150) in MP buffer (20 mM HEPES, pH 7.4, 150 mM NaCl). 20 µl diluted FERRY-RNA mixture (10 nM final in MP buffer) was spotted into a reusable culture well gasket with 3 mm diameter and 1 mm depth (Grace Bio-Labs). MP signals were recorded for 60 s and 120 s at a suitable concentration in order to detect a sufficient set of target particles (>500). Raw MP data were processed in the DiscoverMP software (Refeyn, Oxford, UK).

### Crystallization of Fy-4 and Fy-5

All crystallization experiments were carried out by the sitting-drop method in SWISSCI MRC 2-well crystallization plates at room temperature with a reservoir volume of 50 µl and a drop volume of 3 µl, using a 1:1 mixture of protein and crystallization solution. Initial crystals of Fy-5 were obtained from a 15 mg/ml solution after 4-6 weeks in 0.1 M MES, pH 5.0, 0.8 M Ammonium sulfate. Fy-4 crystals were grown from a 12 mg/ml solution after 3-5 days in 0.1 M MES pH 6.0, 5 % (w/v) PEG 3000 and 30 % (w/v) PEG 200.

### Data collection, structure determination and analysis

Crystals were flash-frozen in liquid nitrogen after a short incubation in a cryo-protecting solution composed of mother-liquor supplemented with 20 % (v/v) glycerol. Data collection was performed at the European Synchrotron Radiation Facility (ESRF) in Grenoble, France under cryogenic conditions at the beamline: ID30A-3. Data were recorded with an EIGER X 4M detector. Diffraction data was processed using XDS (Kabsch, 2010) and the CCP4-implemented program SCALA (Winn et al., 2011). The structures of Fy-4 and Fy-5 were solved by molecular replacement (MR) with CCP4-integrated PHASER (McCoy et al., 2007). APC35852, a member of the DJ superfamily (pdb: 1u9c) was used as search model for Fy-5 and the NADP^+^ bound version of human zeta-crystallin (pdb: 1yb5) was used as search model to solve the structure of Fy-4. The structures were manually built in COOT (Emsley et al., 2004) and refined using PHENIX refine (Adams et al., 2010).

### Hydrogen-Deuterium Exchange Mass Spectrometry

HDX-MS was performed as previously described (Lauer et al., 2019; Mayne et al., 2011; Walters et al., 2012). Proteins (120 µL of 0.5 µM) are diluted 6:4 with 8 M urea, 1% trifluoroacetic acid, passed over an immobilized pepsin column (2.1 mm x 30 mm, ThermoFisher Scientific) in 0.1 % trifluoroacetic acid at 15 °C. Peptides are captured on a reversed-phase C8 cartridge, desalted and separated by a Zorbax 300SB-C18 column (Agilent) at 1 °C using a 5-40 % acetonitrile gradient containing 0.1% formic acid over 10 min and electrosprayed directly into an Orbitrap mass spectrometer (LTQ-Orbitrap XL, ThermoFisher Scientific) with a T-piece split flow setup (1:400). Data were collected in profile mode with source parameters: spray voltage 3.4 kV, capillary voltage 40 V, tube lens 170 V, capillary temperature 170 °C. MS/MS CID fragment ions were detected in centroid mode with an AGC target value of 104. CID fragmentation was 35% normalized collision energy (NCE) for 30 ms at Q of 0.25. HCD fragmentation NCE was 35eV. Peptides were identified using Mascot (Matrix Science) and manually verified to remove ambiguous peptides. For measurement of deuterium uptake, 12 µL of 5 µM protein was diluted in SEC buffer prepared with deuterated solvent. Samples were incubated for varying times at 22 °C followed by the aforementioned digestion, desalting, separation and mass spectrometry steps. The intensity weighted average m/z value of a peptide’s isotopic envelope was compared plus and minus deuteration using the HDX workbench software platform (Pascal et al., 2012). Individual peptides were verified by manual inspection. Data were visualized using Pymol. Deuterium uptake was normalized for back-exchange when necessary, by comparing deuterium uptake to a sample incubated in 6M urea in deuterated buffer for 12 – 18 h at room temperature and processed as indicated above.

### UV-light induced protein-RNA crosslinking

The purified FERRY complex was reconstituted with mrpl41 mRNA in equimolar amounts at 37 °C for 1 h in 20 mM HEPES pH 7.5, 250 mM NaCl, 20 mM KCl, 20 mM MgCl_2_. For reconstitution, the RNA/FERRY complex concentration was adjusted to 400 nM. Aliquots containing 125 pmol of the complex were UV-irradiated (λ = 254 nm) on ice for 10 min in an in-house built crosslinking apparatus following ethanol-precipitation (Kramer et al., 2014). Further sample processing was performed as described with minor modifications (Kramer et al., 2014). Briefly, the protein-RNA pellet was dissolved in 4 M urea, 50 mM Tris/HCl, pH 7.5 by sonication. For RNA digestion, the sample was diluted to 1 M urea with 50 mM Tris/HCl, pH 7.5 and 10 μg RNase A (EN0531, Thermo Fisher Scientific) and 1kU RNase T1 (EN0531, Thermo Fisher Scientific) were added following incubation at 37 °C for 4 h. Proteins were digested over night at 37 °C with trypsin (V5111, Promega) at a 1:20 enzyme to protein mass ratio. Sample cleanup was performed using C18 columns (74-4601, Harvard Apparatus) according to the manufacturers’ instructions and crosslinked peptides were enriched with TiO2 columns (in-house; Titansphere 5 μm; GL Sciences), as described (Kramer et al., 2014). Peptide-(oligo)nucleotides were dried and subjected to LC-ESI-MS/MS.

### Protein-protein crosslinking

For protein-protein crosslinking with EDC, purified FERRY complex was diluted to 400 nM complex concentration using buffer containing 250 mM NaCl, 20 mM KCl, 20 mM MgCl_2_, 20 mM HEPES pH 6.7, 2 mM EDC and 0.5 mM Sulfo-NHS were added, following incubation at 30 °C for 30 min. For quenching, 50 mM β-mercaptoethanol and 20 mM Tris-HCl, pH 7.5 were added. Samples were supplemented with 1 M urea and reduced by addition of 10 mM DTT and 30 min incubation at 37 °C, following alkylation for 30 min at 25 °C with 40 mM iodacetamide (IAA) and quenching of residual IAA by adding another 10 mM of DTT and incubation for 5 min at 37 °C. Protein digestion was accomplished by overnight incubation at 37 °C with trypsin (V5111, Promega) at a 1:20 enzyme to protein mass ratio. C18 columns (74-4601, Harvard Apparatus) were used for peptide clean-up according to the manufacturers’ instructions. Crosslinked peptides were pre-fractionated by high pH reversed-phase chromatography or by peptide size exclusion chromatography.

### High pH reversed-phase chromatography

EDC-crosslinked FERRY peptides were separated on a Xbridge C18 column (Cat. No. 186003128, Waters) using an Agilent 1100 series chromatography system. The system was operated at a flow rate of 60 µL/min with a buffer system comprising: buffer A: 10 mM NH_4_OH; buffer B: 10 mM NH_4_OH pH 10, 80% [v/v] acetonitrile (ACN). The following gradient was used for peptide separation: 5% buffer B (0-7 min), 8-30% buffer B (8-42 min), 30-50% buffer B (43-50 min), 90-95% buffer B (51-56 min), 5% buffer B (57-64 min). For the first 4 min, eluant was collected as flow-through fraction. For the following 48 min, 1-min fractions were collected and reduced to 12 fractions by concatenated pooling. After drying, samples were subjected to LC-MS/MS analysis.

### Peptide size-exclusion chromatography

EDC-crosslinked FERRY peptides were separated on a Superdex Peptide PC3.2/30 column (GE Healthcare) at a flow rate of 50 µL min^−1^. The system was operated in 30% [v/v] CAN, 0.1% [v/v] trifluoroacetic acid (TFA) for 60 min and 1-min fractions were collected. Vacuum-dried fractions were subjected to LC-MS/MS analysis.

### LC-ESI-MS/MS and data analysis

Fractionated EDC-crosslinked peptides or enriched peptide-(oligo)nucleotides were dissolved in 2% [v/v] CAN, 0.05% [v/v] TFA. LC-MS/MS analyses were performed on an Orbitrap Exploris 480 (Thermo Scientific) instrument coupled to a nanoflow liquid chromatography system (Thermo Scientific Dionex Ultimate 3000). Sample separation was performed over 58 min at a flow rate of 300 nL/min using 0.1% [v/v] formic acid (FA) (buffer A) and 80% [v/v] CAN, 0.08% [v/v] FA (buffer B) and a linear gradient from 10% to 45% or to 46% buffer B in 44 min, for peptide-(oligo)nucleotides and EDC-crosslinked peptides, respectively. Eluting peptides were analyzed in positive mode using a data-dependent top 30 acquisition method. Resolution was set to 120,000 (MS1) and 30,000 FWHM (MS2). AGC targets were set to 1e6 (peptide-(oligo)nucleotides) or 3e6 (EDC-crosslinked peptides) for MS1 and 1e5 for MS2. Normalized collision energy was set to 28%, dynamic exclusion to 10 s (peptide- (oligo)nucleotides) or 30 s (EDC-crosslinked peptides), and maximum injection time to 60 ms (peptide-(oligo)nucleotides) or ‘auto’ (EDC-crosslinked peptides) for MS1 and 120 ms for MS2. For peptide-(oligo)nucleotide analyses, charge states 1 and ≥ 8 were excluded; for EDC-crosslinked peptide analyses, charge states 1,2 and ≥ 8 were excluded from fragmentation. Measurements of peptide-(oligo)nucleotides were performed twice for the first technical replicate and once for second and third technical replicate; measurements of EDC-crosslinked peptide fractions were performed once.

Protein-RNA crosslink MS data were analyzed and manually validated using the OpenMS pipeline RNPxl and OpenMS TOPPASViewer (https://www.openms.de/, version 2.6.0) (Kramer et al., 2014). Methionine oxidation was set as variable modification, precursor mass tolerance was set to 6 ppm, maximum missed cleavages to 2, minimum peptide length to 5 amino acids, and maximum number of nucleotides was set to 3. We note that U-H_2_O cannot be distinguished from C-NH_3_ as these RNA adducts have the same monoisotopic masses. EDC crosslink MS raw data files were converted to mgf format using Proteome Discoverer™ software (Thermo Scientific, version 2.1.0.81; signal-to-noise ratio: 1.5; precursor mass: 350– 7000 Da). Crosslink peptide spectra were analysed using pLink software (pFind group, version 1.23) choosing ‘conventional crosslinking (HCD)’ flow type and EDC-DE as linker. Oxidation of methionine was set as variable and carbamidomethylation of cysteine as fixed modification. Precursor mass tolerance was set to 10 ppm, maximum missed cleavages to 2 and minimum peptide length to 5 amino acids. Data were filtered for 1% crosslink spectrum match (CSM) FDR. Downstream analysis and crosslink visualization was performed using xiNET (https://xiview.org/xiNET_website/) (Yang et al., 2012a) and Xlink Analyzer tool (version 1.1.2beta) (Kosinski et al., 2015) in UCSF Chimera (https://www.cgl.ucsf.edu/chimera/, version 1.16).

### Rotary Shadowing

Low-angle metal shadowing and electron microscopy was performed as described previously (Huis in ’t Veld et al., 2016). In brief, freshly purified FERRY complexes were diluted 1:1 with spraying buffer (200 mM ammonium acetate and 60% glycerol) to a concentration of approximately 0.5 μM and air-sprayed onto freshly cleaved mica pieces (V1 quality, Plano GmbH). Specimens were mounted and dried in a MED020 high-vacuum metal coater (Bal-tec). A platinum layer of approximately 1 nm and a 7 nm carbon support layer were subsequently evaporated onto the rotating specimen at angles of 7° and 45°, respectively. Pt/C replicas were released from the mica on water, captured by freshly glow-discharged 400-mesh Pd/Cu grids (Plano GmbH), and visualized using a LaB_6_ equipped JEM-1400 transmission electron microscope (JEOL) operated at 120 kV. Images were recorded at a nominal magnification of 60,000x on a 4k x 4k CCD camera F416 (TVIPS), resulting in 0.189 nm per pixel. Particles with discernible coiled-coil extensions were manually selected.

### Sample vitrification

For sample preparation in cryo-EM, 3.5 µL of purified FERRY complex at a concentration of 0.7 mg/mL was applied to freshly glow-discharged UltrAuFoil 1.2/1.3 grids (Quantifoil), automatically blotted for 3 s and plunged into liquid ethane using the a Vitrobot (Thermo Fisher Scientific), operated at 100 % humidity and 13 °C. Individual grid quality was screened prior to data collecting using a Talos Arctica transmission electron microscope (TEM, Thermo Fisher Scientific), operated at 200 kV. Prior to data collection, grids were stored in liquid nitrogen.

### Cryo-EM data acquisition

Cryo-EM data set of the FERRY complex was collected on a Titan Krios TEM (Thermo Fisher Scientific), equipped with an C_s_-Corrector and in-column energy filter, operated at an acceleration voltage of 300 kV. Micrographs were recorded on a K2 direct electron detector (Gatan) with a final pixel size of 1.08 Å in counting mode. A total of 40 frames (with 375 ms and 1.895 e^-^/Å^2^ each) was recorded during each exposure, resulting in a total exposure time of 15 s and an overall electron dose of 75.8 e^-^/Å^2^. Automated data collection was done with the help of the software EPU (Thermo Fisher Scientific) and monitored in real time using TranSPHIRE (Stabrin et al., 2020). A total of 1879 micrographs was collected with a defocus range between −1.6 µm and −2.8 µm and an energy filter width of 20 eV.

### Image processing and 3-D reconstruction

Initially, micrographs were inspected visually to discard images with high drift and ice-contamination. Using MotionCor2, operated in 3 x 3 patch mode, individual frames were aligned and summed (Zheng et al., 2017). In this step, unweighted and dose-weighted full-dose images were calculated. Image processing was performed with the SPHIRE software package (Table S1) (Moriya et al., 2017). Values for the defocus and astigmatism of unweighted full-dose images were determined using CTFFIND4 (Rohou and Grigorieff, 2015). A flowchart of the image processing strategy is described in Fig. S2. First, particles were automatically selected based on a trained model with the help of crYOLO (Wagner and Raunser, 2020; Wagner et al., 2019). After extraction of the particles with a window size of 264 x 264, the resulting stack was further classified using the iterative and stable alignment and clustering (ISAC) algorithm, implemented in SPHIRE. This yielded a stack of 18.5 k particles of dose-weighted drift-corrected particles. Based on a subset of class averages produced by ISAC, the *ab initio* 3D structure determination program RVIPER in SPHIRE calculated an initial intermediate resolution 3D structure that served as reference in the subsequent 3D refinement (MERIDIEN). This 3D refinement step, in which C2 symmetry was imposed, yielded a 5.9 Å map of the core of the FERRY complex, estimated by the ‘gold standard’ 0.143 criterion of the Fourier shell correlation (FSC). Based on the obtained 3D parameters, particles were re-centered, followed by re-extraction, resulting in 18.3 k particles. 3D refinements with C2 symmetry and without applying symmetry yielded 3D reconstruction with resolutions of 4.6 and 6.2 Å, respectively. In the next step, iterative cycles of Bayesian particle polishing in RELION (Scheres, 2012) and 3D refinement in SPHIRE (Moriya et al., 2017) were performed. The particles of this ‘polished’ stack were further subjected to another round of 3D refinement in SPHIRE with imposed C2 symmetry. Here, the real space filtering according to signal-to-noise ratio (SNR) algorithm, named SIDESPLITTER, was applied to reduce overfitting (Ramlaul et al., 2020). This resulted in a final 4.0 Å electron density map of the FERRY complex.

In general, the global resolution of maps was calculated between two independently refined half maps at the 0.143 criterion. Local resolutions were calculated with LOCALRES in SPHIRE. EM density maps were either filtered according to global resolution or using the local de-noising filter L-AFTER (Ramlaul et al., 2019). In the latter case, which is also based on half maps, features with more signal than noise are better recovered.

### Model building, refinement and validation

To build the model for the (Fy-4)_2_(Fy-2)_2_(Fy-5)_4_ core of the FERRY complex, the obtained crystal structures of Fy-4 and Fy-5 were initially fitted into the corresponding density using the rigid body fitting tool in Chimera (Pettersen et al., 2004). trRosetta, a *de novo* protein structure prediction algorithm that is based on direct energy minimization with restrained Rosetta, was used to obtain initial models for Fy-2 (Yang et al., 2020). The predicted model for the 6-helix bundle domain, containing residues 246 to 498, that matched our experimental density best was subsequently fitted similarly to Fy-4 and Fy-5 using rigid body fit. Manual model building for the regions N- and C-terminal 6-helix bundle, which comprise residues 218 – 245 and 499 – 552, respectively, was further guided by secondary structure predictions of individual trRosetta runs for these regions, which include the vertical helix as well as the beginning of the two terminal coiled-coils of Fy-2. With the resulting combined model, containing residues 2-349, 218-552 and 8-217 of Fy-4, Fy-2 and Fy-5, respectively, a restrained refinement in PHENIX was performed (Liebschner et al., 2019). In the next step, the model was further refined using a combination of manual building in COOT and real-space refinement in PHENIX (Emsley et al., 2004; Liebschner et al., 2019). Geometries of the final model were either obtained from PHENIX or calculated using Molprobity (http://molprobity.biochem.duke.edu). Refinement and model building statistics are summarized in Table S1.

### Hierarchical classification of 2-D classes

Separate hierarchical classifications were run for classes selected from ISAC (Yang et al., 2012b). Aligned particles from each class were generated using the SPHIRE program *sp_eval_isac.py* (http://sphire.mpg.de/wiki/doku.php?id=pipeline:utilities:sp_eval_isac). Particles for each class were then subjected to multivariate data analysis and hierarchical classification in SPIDER (Frank et al., 1996; Shaikh et al., 2008). Binary masks for correspondence analysis were drawn manually onto class averages using *e2display.py* from EMAN2 (Tang et al., 2007), and then thresholded in SPIDER using the operation ‘TH M’. Hierarchical classification using Ward’s method was performed using SPIDER operation ‘CL HC’. Class averages were visualized by the Python script *binarytree.py* which uses SPIDER’s SPIRE libraries (Baxter et al., 2007). SPIDER procedures can be found at the SPIDER web site (https://github.com/spider-em/SPIDER/tree/master/docs/techs/MSA).

### Structure analysis and visualization

UCSF Chimera was used for structure analysis, visualization and figure preparation (Pettersen et al., 2004). The angular distribution plots as well as beautified 2-D class averages were calculated in SPHIRE (Moriya et al., 2017).

## SI Movies

**Movie S1. Flexibility of the C-terminal coiled-coil domain of Fy-2 in the FERRY complex.**

This movie highlights multiple orientations that the C-terminal coiled-coil region of Fy-2 can adopt relative to the core of the FERRY complex. Related to Fig. 4.

**Movie S2. Flexibility of the N-terminal coiled-coil domain of Fy-2 in the FERRY complex.**

This movie highlights multiple orientations that the N-terminal coiled-coil domain of Fy-2 can adopt relative to the FERRY core. Related to Fig. 4.

## SI Files

**File S1. Crosslinking mass spectrometry of FERRY with mRNA. Related to** Fig. 5.

**File S2. Crosslinking mass spectrometry of FERRY with mRNA. Related to Fig. S5.**

## Acknowledgments

We thank O. Hofnagel and D. Prumbaum for assistance with EM data collection, M. Stabrin for lively discussions regarding image processing and M. Raabe for help with MS of crosslinked samples. We also acknowledge R. Goody and A. Musacchio for valuable feedback regarding the manuscript. We thank the members of the cluster of excellence “Physics of Life” (Deutsche Forschungsgemeinschaft under Germany’s Excellence Strategy—EXC-2068–390729961— Cluster of Excellence Physics of Life of Technische Universität Dresden) for stimulating discussion. This research was financially supported by the Deutsche Forschungsgemeinschaft (DFG, German Research Foundation) - Project Number 112927078 - TRR 83 (to M.Z.), and the Max Planck Society (to S.R., M.Z. and H.U.). J.S.S. was funded by the Deutsche Forschungsgemeinschaft (DFG, German Research Foundation) - Project Number 112927078 - TRR 83.

## Author contributions

J.S.S., S.R. and M.Z. designed the project. J.S.S. provided the protein complex and performed X-ray studies. J.S.S. and J.L. performed the hydrogen-deuterium exchange mass spectrometry experiments, L.M.W. and H.U. performed and analyzed crosslinking-MS experiments and P.J.H. performed low-angle platinum shadowing experiments. D.Q. prepared specimens, recorded, analyzed and processed the electron microscopy data and prepared figures. T.R.S. performed hierarchical classification analysis. D.Q. and B.U.K. built the atomic models. S.R. managed the project. D.Q. and S.R. wrote the manuscript with input from all authors.

## Data availability

The cryo-EM density map of the FERRY complex is deposited into the Electron Microscopy Data Bank with the accession number EMD-12273. Model coordinates for the FERRY complex, the X-ray structures of Fy-4 and Fy-5 are available with the PDB entry IDs 7ND2, 8A3O and 8A3P, respectively. The mass spectrometry proteomics data have been deposited to the ProteomeXchange Consortium via the PRIDE partner repository with the dataset identifier PXD034875.

