## Supplementary material for "Structure of the human FERRY Rab5 effector complex": Supplemtal Figures and Tables

#### SI Figures

A) Fy-5

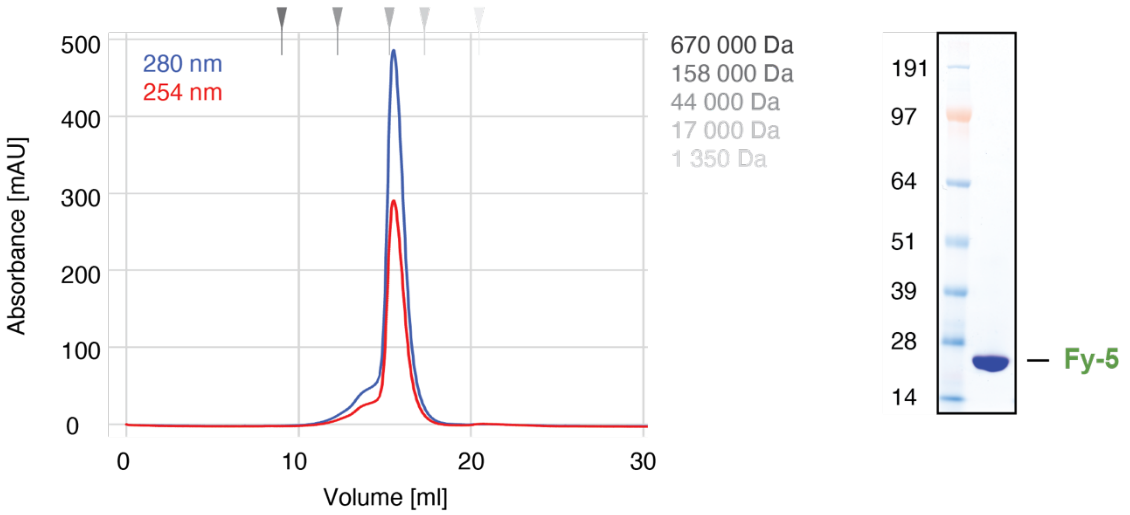


B) Fy-4

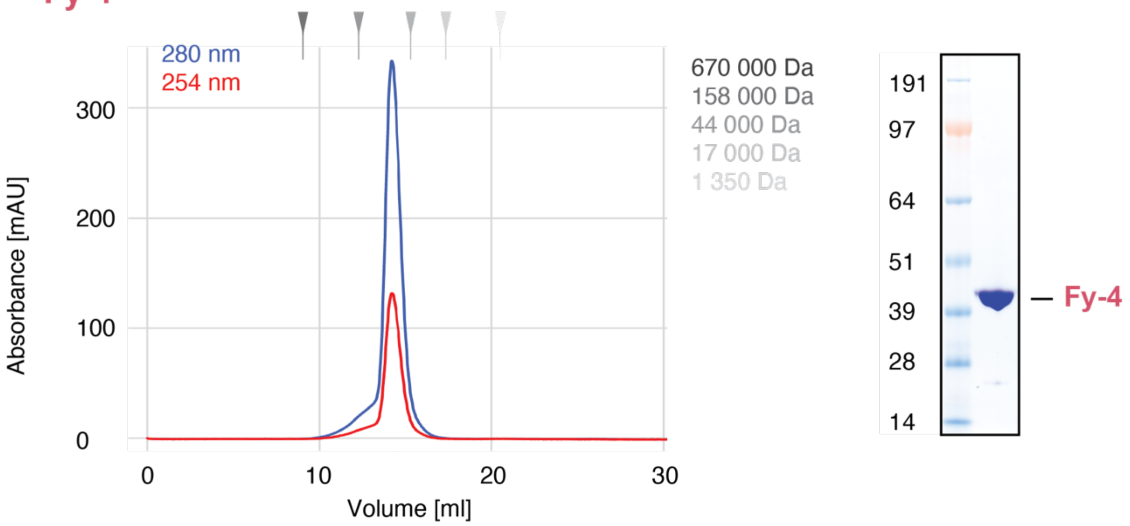


C) FERRY complex

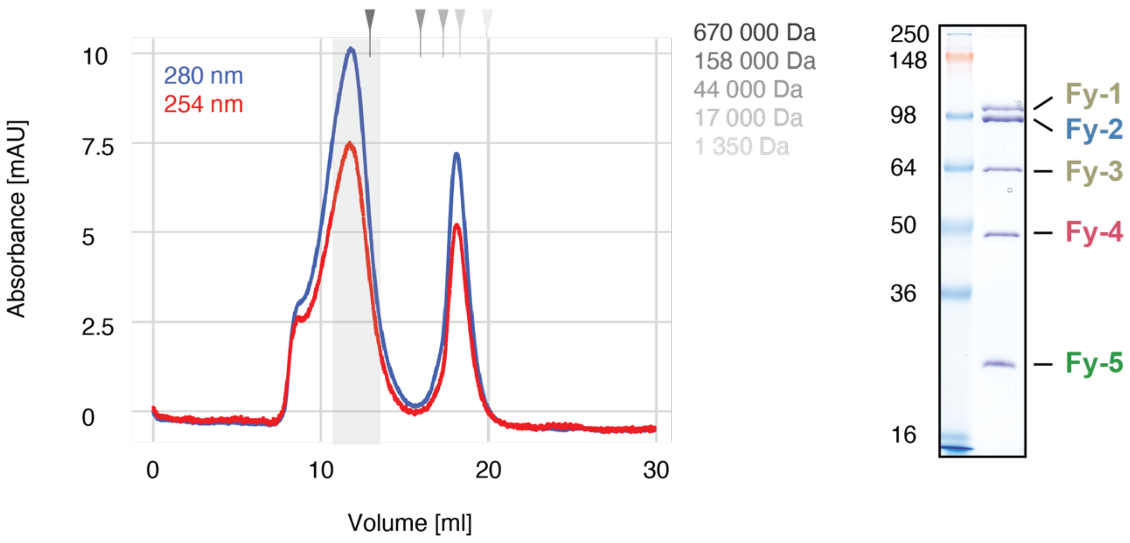

#### Figure S1. Purification of the FERRY complex.

(**A-C**) Size exclusion chromatography profiles (left) and corresponding Coomassie-stained SDS-PAGE (right) of purified Fy-5 (**A**), Fy-4 (**B**) and the FERRY complex (**C**). Positions of protein standards are indicated by marks. Grey box highlights fractions used for subsequent analysis.
Related to Fig. 1.

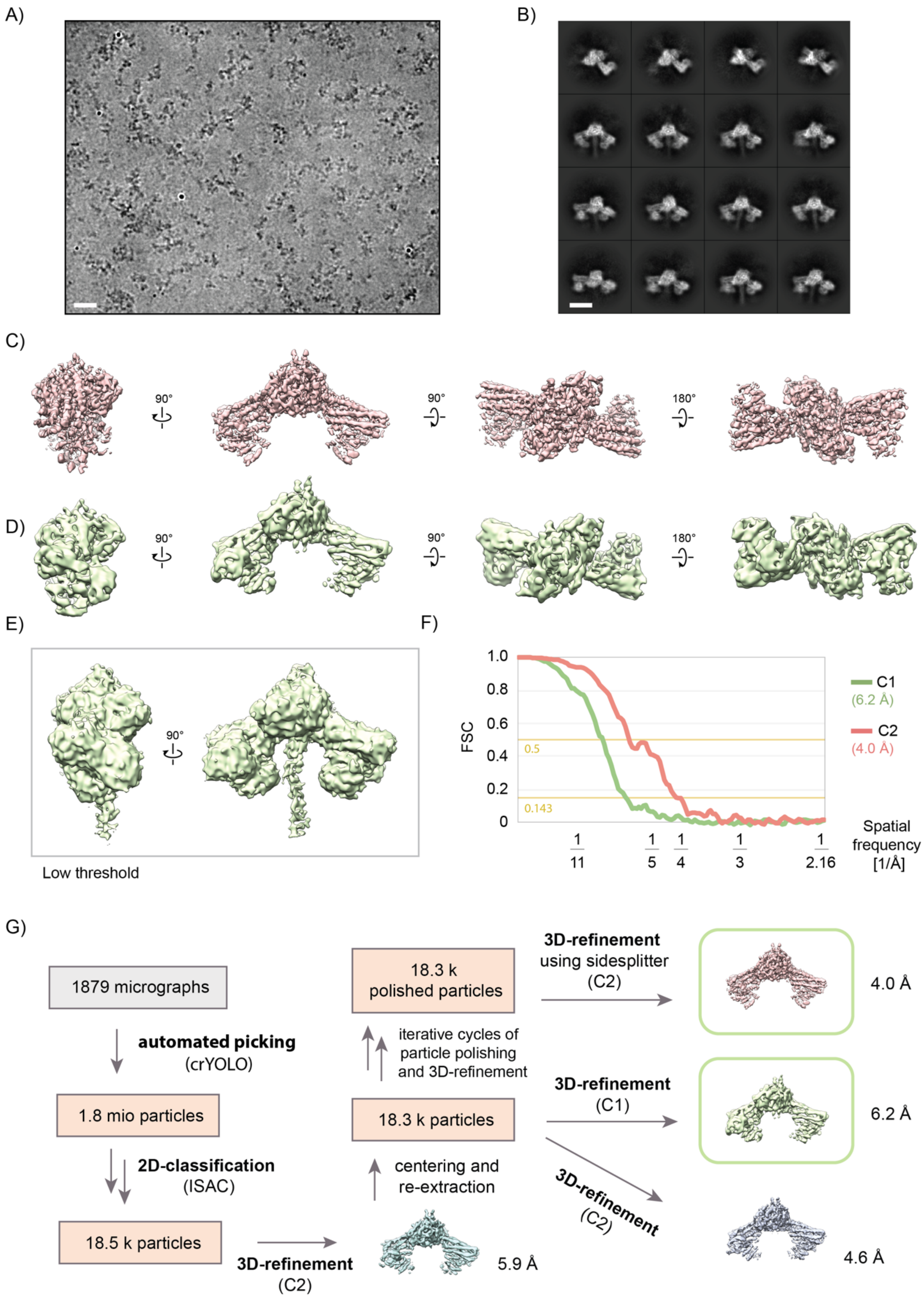


**Figure S2. Cryo-EM of the FERRY complex.**

(**A**) Representative JANNI-filtered digital micrograph area of vitrified FERRY complex. Scale bar: 20 nm.
(**B**) Representative 2-D class averages corresponding to (A). Scale bar: 10 nm.

(**C-D**) Rotated views of the 3-D reconstruction of the core of the FERRY complex with C2 symmetry applied (C) and without applying symmetry (D).
(**E**) Reconstruction corresponding to (D) is shown at a lower threshold to highlight the density corresponding to the N-terminal coiled-coil of Fy-2.
(**F**) FSC curves between two independently refined half maps of the FERRY core with C2 symmetry applied (C2, red) and without applying symmetry (C1, green).
(**G**) Flowchart of image processing strategy. After automated particle selection with crYOLO, 2-D classification was performed using the ISAC algorithm implemented in SPHIRE, yielding a total of 18.5 k particles. Initial 3-D refinement (MERIDIEN in SPHIRE) was followed by centering and subsequent re-extraction. The resulting particle stack was used to calculate two 3-D reconstructions, with C-2 symmetry applied and without symmetry. In the next step, the particle stack was further polished by iterative cycles of particle polishing and 3-D refinement. Polished particles were subjected to another round of 3-D refinement in which the real space filtering algorithm SIDESPLITTER was applied, resulting in a 4.0 Å 3-D reconstruction of the core of the FERRY complex.
Related to Fig. 1.

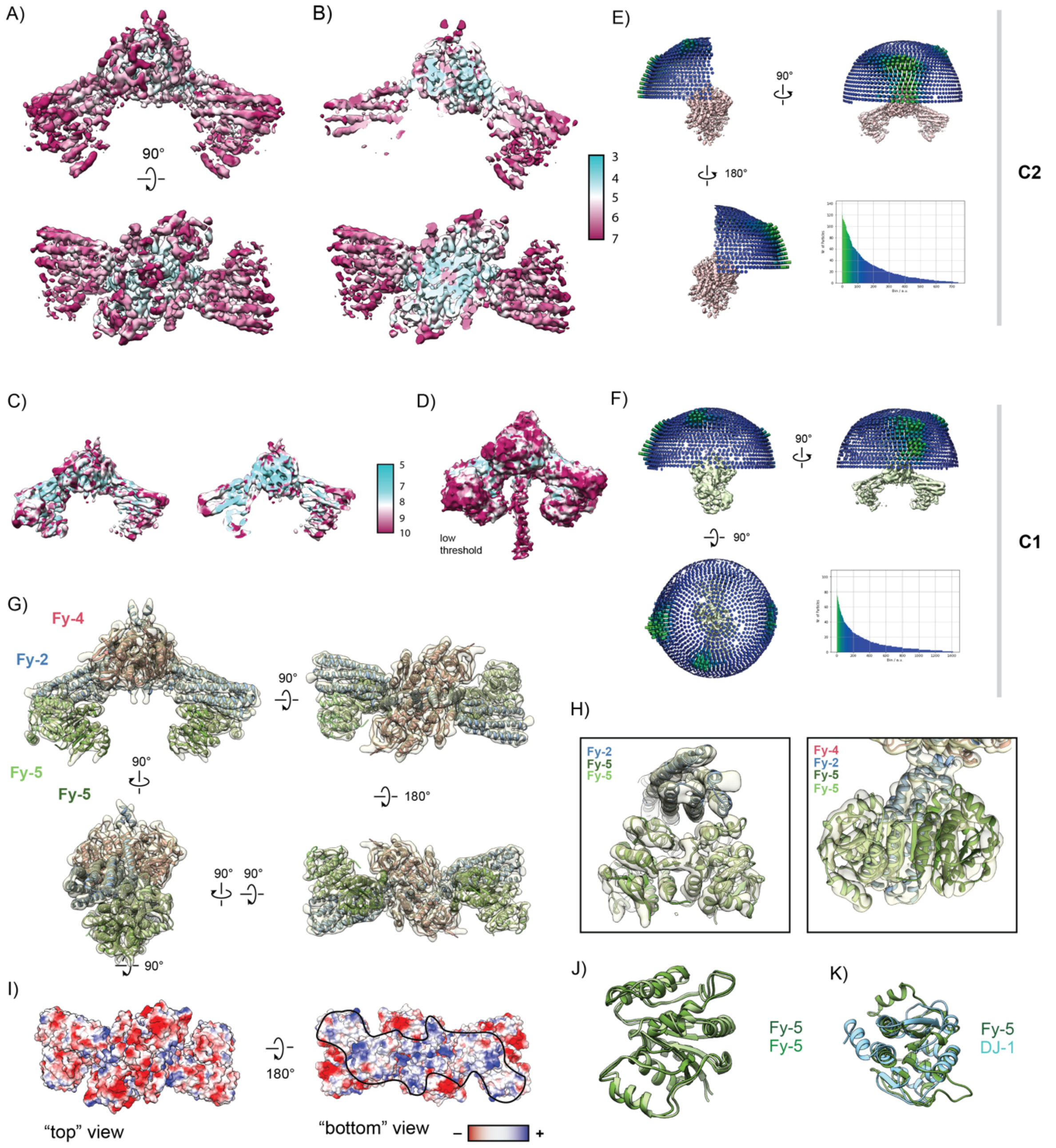


**Figure S3. Local resolution and 3-D orientation plots of the FERRY complex.**

(**A**) Rotated views of the 3-D reconstruction of the FERRY complex core with C2 symmetry applied, colored by local resolution.
(**B**) Cross-sections corresponding to (A). Color key of local resolution in Å is provided on the right.
(**C**) 3-D reconstruction of the core of the FERRY complex without applying symmetry (left) and corresponding cross-section (right), colored by local resolution. Color key of local resolution in Å is provided next to it.
(**D**) Same 3-D reconstruction as in (C) is shown at lower threshold to illustrate the density corresponding to the N-terminal coiled-coil domain of Fy-2. Same color code as in (C).
(**E-F**) Rotated views of the 3-D angular distribution plots for the reconstruction with applied C2 symmetry (E) and without symmetry (F). The relative height of bars represents the number of particles. Corresponding 2-D histogram is provided on the lower right.
(**G**) Rotated views of the L-AFTER-filtered 3-D reconstruction of the FERRY complex core with the corresponding atomic model fitted inside. Fy-2 – blue; Fy-4 – red; Fy-5 (distal) – light green; Fy-5 (proximal) – dark green.

(**H**) Close-ups of the “arm” region, consisting of the 6-helix bundle domain of Fy-2 and the Fy-5 dimer. The left panel shows a view perpendicular to the central axis of the 6-helix bundle domain, whereas in the right panel a view along the C2 axis of the Fy-5 dimer is displayed.
(**I**) Top- and bottom view of the electrostatic surface of the FERRY complex core. Interestingly, in contrast to the “top” side, the “bottom” side is lined with positively charged residues. Boundaries of patches are indicated by black line.
(**J**) Superposition of distal (dark green) and proximal (light green) Fy-5 shows that both proteins are almost identical at the backbone level (RMSD – 0.943 Å).
(**K**) Superposition of distal Fy-5 (green) and the recessive parkinsonism protein DJ-1 (cyan) highlights a common core fold shared by both proteins (RMSD – 1.012 Å). Differences are localized primarily to the periphery of the protein.
Related to Fig. 1.

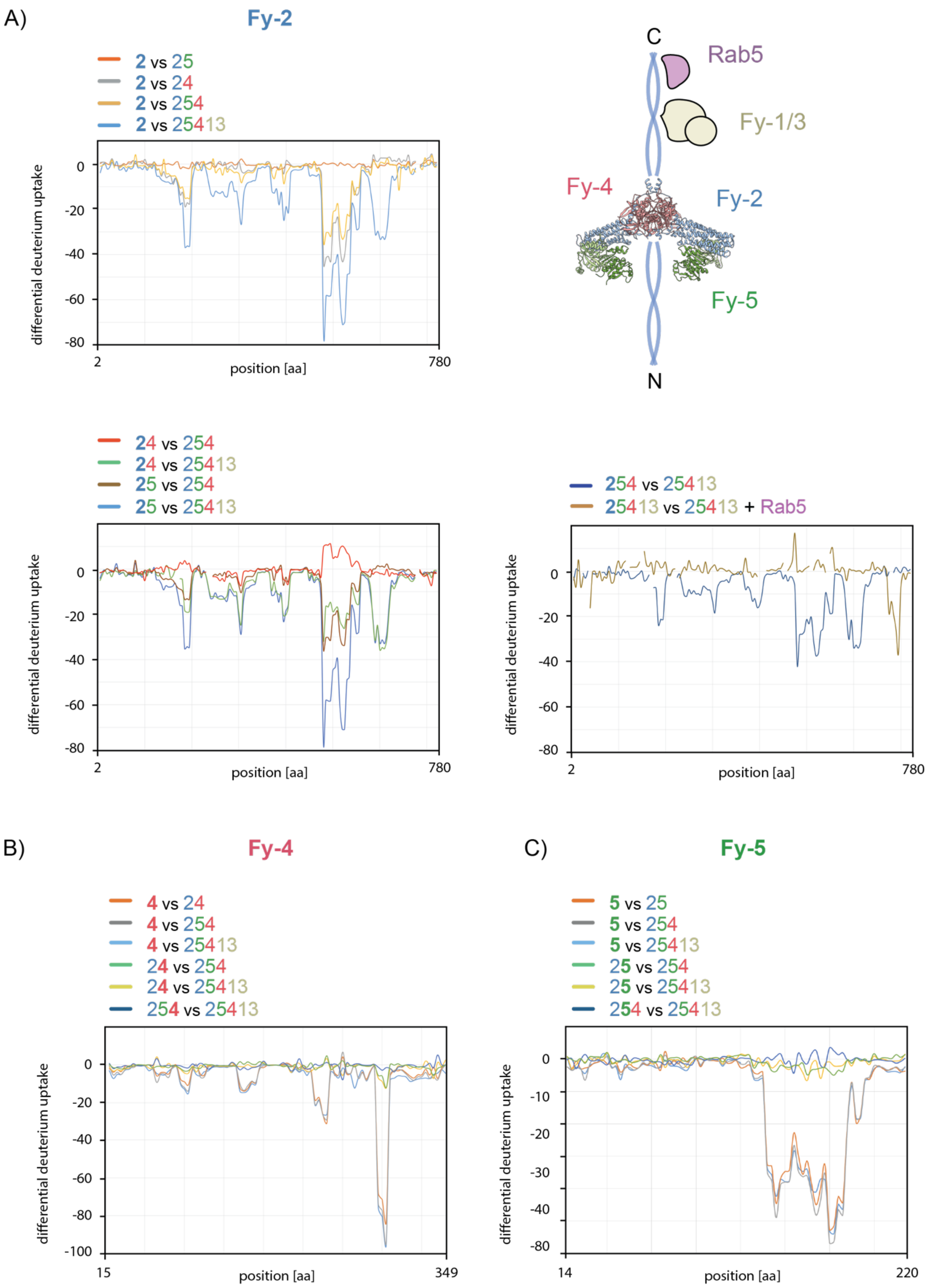

A)
Fy-2
2 vs 25
2 vs 24
2 vs 254
2 vs 25413
differential deuterium uptake
0
-20
-40
-60
-80
2
position [aa]
780
C
Rab5
Fy-1/3
Fy-4
Fy-2
Fy-5
N
24 vs 254
24 vs 25413
25 vs 254
25 vs 25413
254 vs 25413
25413 vs 25413 + Rab5
B)
Fy-4
4 vs 24
4 vs 254
4 vs 25413
24 vs 254
24 vs 25413
254 vs 25413
-100
15
349
C)
Fy-5
5 vs 25
5 vs 254
5 vs 25413
25 vs 254
25 vs 25413
254 vs 25413
-10
-30
14
220

**Figure S4. Hydrogen-deuterium exchange mass spectrometry delineates the binding interfaces of the FERRY complex.**

(**A**) In the upper left panel and lower two panels, differential deuterium uptake (DDU) of Fy-2 is plotted against detected peptides (as peptides can overlap, they are approximated by the amino acid sequence). DDU profiles are color coded with the respective legend shown next to the plot. Peptides of the subunit displayed in bold are plotted, while interruptions in the profiles being caused by absence of detected peptides. The schematic representation depicted in the upper right panel explains the abbreviations used in the profile legend.
(**B, C**) Similar DDU profiles as described in (A) for Fy-4 (B) and Fy-5 (C).
Related to Fig. 2-4.

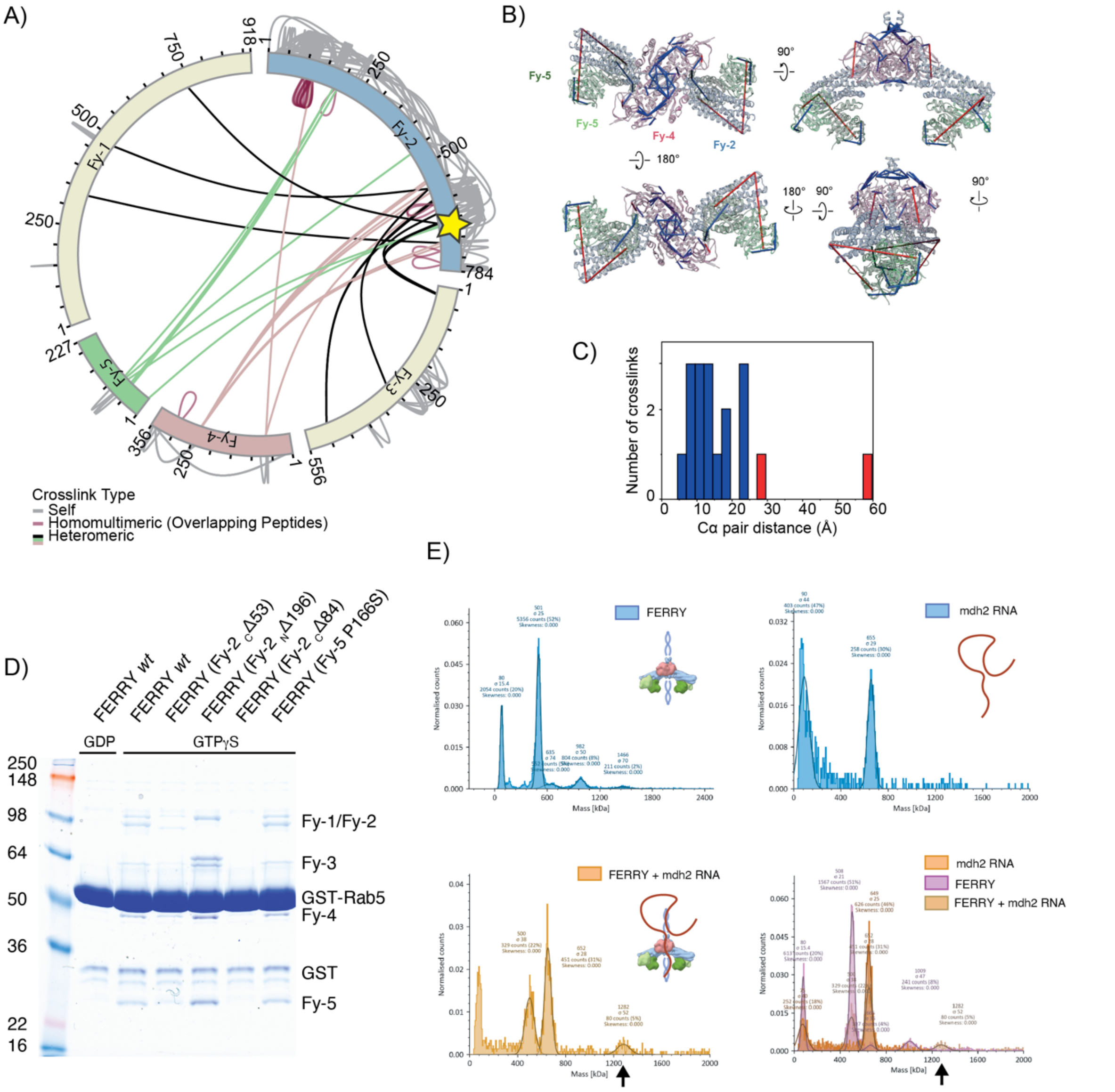


### Figure S5. Cross-linking MS and mass photometry of FERRY and Rab5 pulldown assay.

(**A**) FERRY complex crosslink network identified by EDC-crosslinking mass spectrometry. Inter- (black, green, pink), intra-protein crosslinks (grey) and loop links (red) identified at a 1% FDR are shown as lines. Fy-1/3 binding site, identified by HD-X mass spectrometry is indicated by yellow star.
(**B**) EDC crosslinks mapped to the structure of Fy2-Fy4-Fy5 complex. Lines connect Cα atoms of crosslinked amino acid residues. Blue and red lines represent crosslinks showing distances below and above 25 Å, respectively.
(**C**) Histogram of EDC Fy2-Fy4-Fy5 crosslink distances between crosslinked Cα atoms. 89% of crosslinks meet the distance restraint of 25 Å. 11% of crosslinks violate the distance restraint which might be effect of complex structure dynamics or experimental errors.
(**D**) GST-Rab5 pulldown assay with the FERRY complex in the presence of GDP or GTPγS. With GTPγS, different Fy-2 truncation variants as well as the Fy-5 P166S mutant were tested.
(**E**) Mass photometry measurements of the FERRY complex alone (upper left panel), *mdh2* mRNA alone (upper right panel), and when FERRY and mdh2 mRNA were mixed (lower left panel). Bottom right panel shows the overlap of the three individual measurements. Note the new emerging peak at 1282 kDa upon interaction of FERRY and RNA, indicated by a black arrow.
Related to Fig. 4. See also File S2.

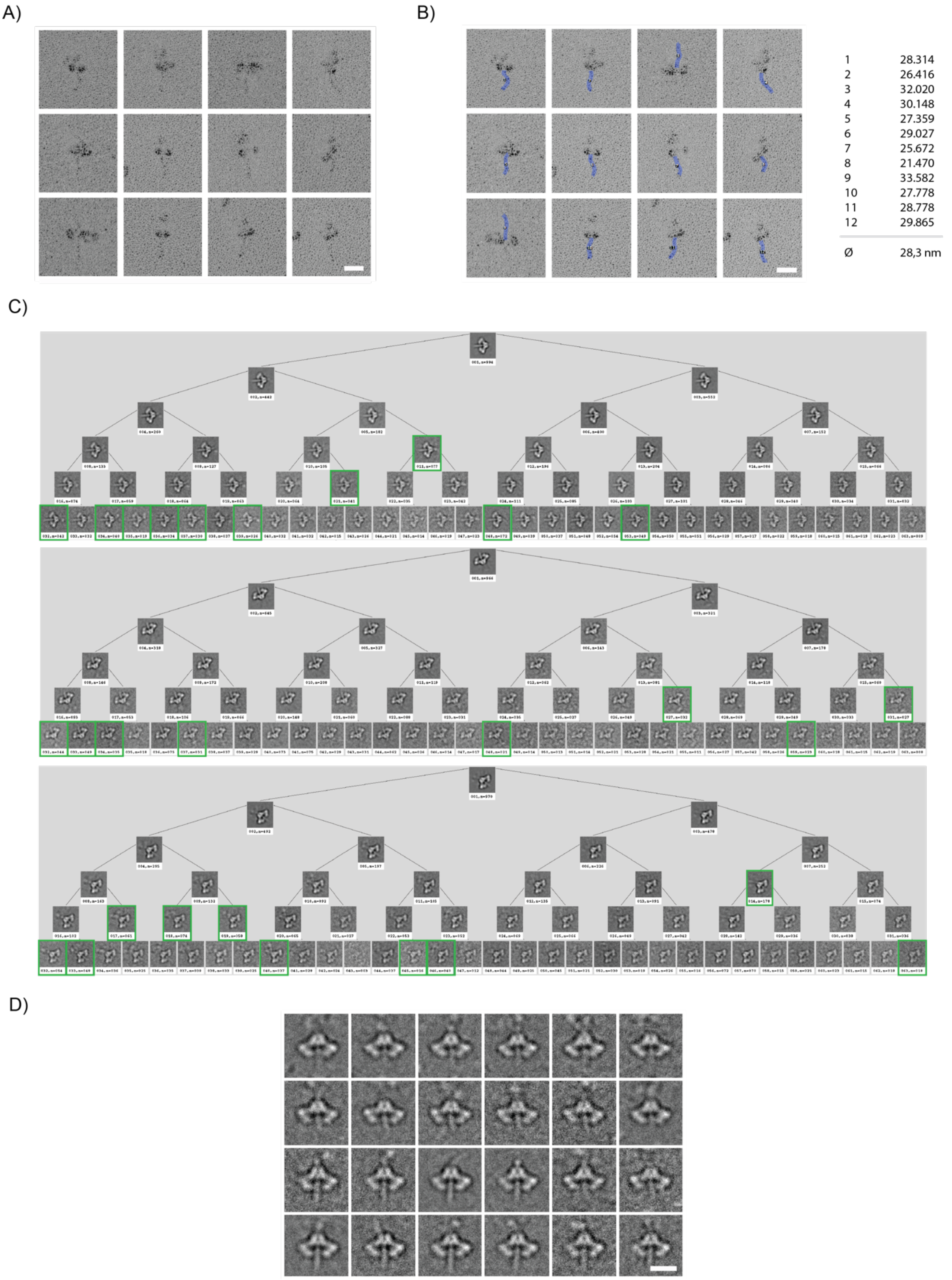


**Figure S6. Rotary shadowing EM and hierarchical classification of 2D cryo-EM classes of the FERRY complex.**

(**A**) Selected examples of FERRY complex visualized by electron microscopy after glycerol spraying and low-angle platinum shadowing. Scale bar: 20 nm.
(**B**) The longest visible coiled-coil domain of the particles shown in (A) was measured, resulting in an average length of 28 nm. Scale bar: 20 nm.

(**C**) Cryo-EM 2D classes of FERRY selected from ISAC were run through correspondence analysis in SPIDER (operation 'CA S') and hierarchical classification (operation 'CL HC'). Results were visualized using SPIRE's *binarytree.py*. Classes highlighted in green show density corresponding to the C-terminal coiled-coil domain of Fy-2.
(**D**) Selection of cryo-EM 2D classes showing density corresponding to the C-terminal coiled-coil domain of Fy-2. Scale bar: 10 nm.
Related to Fig. 4.

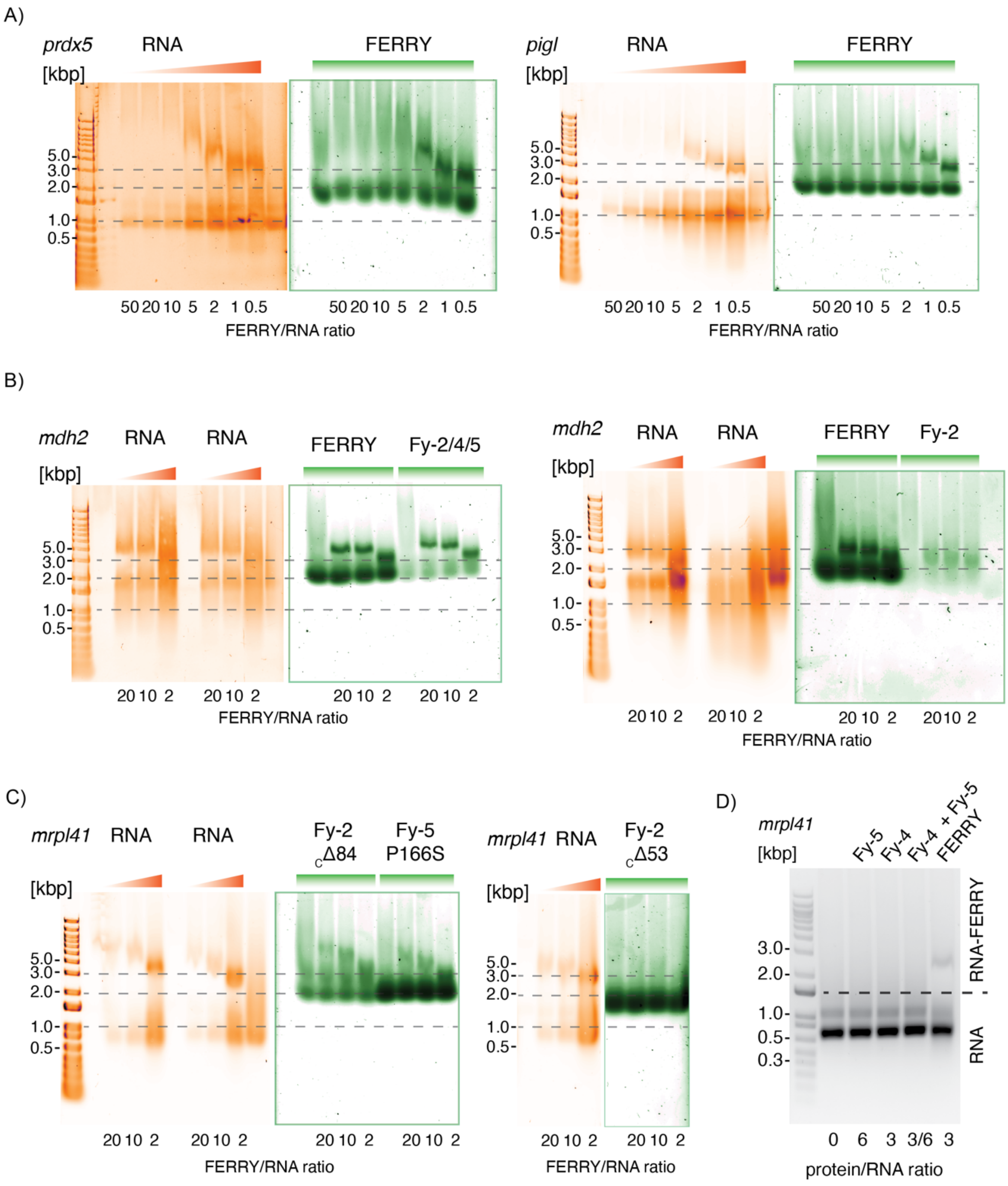


**Figure S7. Electrophoretic mobility shift assays of FERRY and mRNA.**

(**A**) Electrophoretic mobility shift assays (EMSAs) showing the interaction between the FERRY complex and *prdx5* and *pigl* mRNAs in the left and right panel, respectively. RNA is stained with SYBR Gold (orange, left) and protein with Sypro Red (green, right). Note that with decreasing ratios of FERRY:RNA a shift towards higher mobility of the FERRY-RNA complex is observed.
(**B**) EMSAs showing that binding of the 3-subunit Fy-2/4/5 complex to *mdh2* mRNA is comparable to wt FERRY (left panel), whereas Fy-2 alone is not sufficient for mRNA binding (right panel).
(**C**) EMSAs of FERRY with *mrpl41* mRNA and either Fy-2 replaced by C-terminal truncation variants or Fy-5 by mutated Fy-5 bearing the P166S mutation.
(**D**) EMSAs of *mrpl41* mRNA with either Fy-4, Fy-5, Fy-4/Fy-5 or the FERRY complex. Only fully assembled FERRY is capable of binding mRNA.
Related to Fig. 5 and 6.

**SI Tables**

Table S1. Data collection, refinement and model building statistics of FERRY cryo-EM structures. Related to Fig. 1.

| **Microscopy and cryo-EM** | C2 | C1 |
|---|---|---|
| Microscope | Titan Krios | |
| Voltage [kV] | 300 | |
| Camera | K2 Summit | |
| Defocus range [µm] | -1.6 to -2.8 | |
| Pixel size [Å] | 1.08 | |
| Total electron dose [$e^-/Å^2$] | 75.8 | |
| Exposure time [s] | 15 | |
| Frames per movie | 40 | |
| Number of micrographs | 1879 | |
| Number of particles in final reconstruction | 18,3 k | |
| Map resolution [Å] | 4.0 | 6.2 |
| **Model statistics** | | |
| Bond RMSD [Å] | 0.011 | |
| Angle RMSD [°] | 1.66 | |
| Rotamer outliers [%] | 0.21 | |
| Ramachandran - favored [%] | 94.63 | |
| Ramachandran - allowed [%] | 5.05 | |
| Ramachandran - outliers [%] | 0.32 | |
| Molprobity score | 1.96 | |
| EMRinger score | 1.24 | |

Table S2. Data collection, refinement and model building statistics of X-ray structures. Related to Fig. 1.

| **Data collection** | Fy-5 | Fy-4 |
|---|---|---|
| Space group | C222(1) | P2 |
| a, b, c, [Å] | 94.62<br>99.57<br>62.12 | 86.42<br>45.27<br>93.96 |
| $\alpha$, $\beta$, $\gamma$ [Å] | 90<br>90<br>90 | 90<br>109.93<br>90 |
| Energy [keV] | 12.81 | 12.81 |
| Resolution [Å] | 47.31 - 2.7 | 45.92 – 2.9 |
| Inner shell | 2.85 - 2.6 | 3.06 – 2.9 |
| $R_{merge}$ | 0.088 (0.603) | 0.056 (0.209) |
| I/$\sigma$ (I) | 13.4 (3.2) | 17.9 (6.2) |
| Completeness [%] | 99.6 (99.9) | 99.1 (99.7) |
| Redundancy | 9.6 (10.1) | 4.9 (5.0) |
| $CC_{1/2}$ | 0.997 (0.970) | 0.996 (0.974) |
| **Refinement** | Fy-5 | Fy-4 |
| Resolution [Å] | 2.7 | 2.9 |
| No. reflections | 8324 | 15371 |
| $R_{work}/R_{free}$ | 22.0/26.7 | 23.7/28.5 |
| **No. Atoms** | Fy-5 | Fy-4 |
| Protein | 1426 | 4954 |
| Ligand | 0 | 0 |
| Water | 0 | 21 |
| **R.m.s deviations** | Fy-5 | Fy-4 |
| Bond lengths [Å] | 0.006 | 0.004 |
| Bond angles [°] | 0.894 | 0.709 |
| **Ramachandran** | Fy-5 | Fy-4 |
| Ramachandran - favored [%] | 95.8 | 95.0 |
| Ramachandran - allowed [%] | 4.2 | 4.7 |
| Ramachandran - outliers [%] | 0 | 0.3 |

#### SI Movies

**Movie S1. Flexibility of the C-terminal coiled-coil domain of Fy-2 in the FERRY complex.**

This movie highlights multiple orientations that the C-terminal coiled-coil region of Fy-2 can adopt relative to the core of the FERRY complex. Related to Fig. 4.

**Movie S2. Flexibility of the N-terminal coiled-coil domain of Fy-2 in the FERRY complex.**

This movie highlights multiple orientations that the N-terminal coiled-coil domain of Fy-2 can adopt relative to the FERRY core. Related to Fig. 4.

#### SI Files

**File S1. Crosslinking mass spectrometry of FERRY with mRNA.** Related to Fig. 5.
